# Empirically Constrained Network Models for Contrast-dependent Modulation of Gamma Rhythm in V1

**DOI:** 10.1101/729707

**Authors:** Margarita Zachariou, Mark Roberts, Eric Lowet, Peter De Weerd, Avgis Hadjipapas

## Abstract

Here we present experimentally constrained computational models of gamma rhythm and use these to investigate gamma oscillation instability. To this end, we extracted empirical constraints for PING (Pyramidal Interneuron Network Gamma) models from monkey single-unit and LFP responses recorded during contrast variation. These constraints implied weak rather than strong PING, connectivity between excitatory (E) and inhibitory (I) cells within specific bounds, and input strength variations that modulated E but not I cells. Constrained models showed valid behaviours, including gamma frequency increases with contrast and power saturation or decay at high contrasts. The route to gamma instability involved increased heterogeneity of E cells with increasing input triggering a breakdown of I cell pacemaker function. We illustrate the model’s capacity to resolve disputes in the literature. Our work is relevant for the range of cognitive operations to which gamma oscillations contribute and could serve as a basis for future, more complex models.

## Introduction

Gamma oscillations are present throughout the cortex (Bastos et al., 2015; Bosman, Lansink, & Pennartz, 2014) and are thought to play an important role in neural communication and a range of cognitive operations (Bosman et al., 2014; Engel, Fries, & Singer, 2001; Fries, 2009, 2015; Singer, 1995). Improving the understanding of gamma generating mechanisms has therefore been a central question in neuroscience in the last decades. Gamma oscillation frequency changes strongly and monotonically with visual contrast that increases the firing rate of visual neural pathways (Hadjipapas, Lowet, Roberts, Peter, & De Weerd, 2015; Ray & Maunsell, 2010; Roberts et al., 2013). Gamma power varies nonmonotonically with contrast and shows saturation or power decay at high contrasts (Hadjipapas et al., 2015; Roberts et al., 2013), which suggests instability in the underlying network oscillation dynamics. Such a dissociation between frequency and power (see also (Jia, Xing, & Kohn, 2013)) and the possible oscillation instability at high contrasts provide interesting entry points onto the network mechanisms generating gamma oscillation under different input firing conditions. The goal of the present paper was to generate a model that could simultaneously display multiple empirically observed features, including the frequency increase as well as power saturation/decay with contrast (input strength). To that aim, we constrained a gamma generating network model by characteristics of LFP gamma oscillations as well as spiking data measured as a function of contrast in monkeys.

We focused on so-called Pyramidal Interneuron Network Gamma (PING) models, which have provided a good insight into the basic mechanisms of gamma in V1 and other cortical areas. In the PING model, the excitatory cells (E cells) are the drivers of the gamma rhythm (Tiesinga & Sejnowski, 2009). When activated, they stimulate the inhibitory cells (I cells), which in turn provide inhibitory feedback to the E cells, which restart firing once the decay of inhibition is overcome by on-going excitation. In the so-called “strong PING” mechanism (Börgers & Kopell, 2005; Tiesinga & Sejnowski, 2009) both E and I cells spike in a phase-locked manner at a specific phase in the gamma cycle, with E spikes locked to a slightly earlier phase than I spikes. In so-called “weak” PING models, E cells fire irregularly and at a much lower frequency compared to the LFP population gamma rhythm, with only a sparse random set participating in each cycle of gamma (Börgers & Kopell, 2005; Lee & Jones, 2013; Wang, 2010; Whittington, Traub, Kopell, Ermentrout, & Buhl, 2000). I cells, as in the strong PING model, fire at a rate close to the LFP gamma frequency. In another class of network models referred to as the Interneuron Network Gamma (ING) model, the I cell population inhibits itself, and generates oscillatory firing at a gamma frequency determined by the time constant that regulates the decay of inhibition (Tiesinga & Sejnowski, 2009). E cells may be passively entrained by the gamma rhythm imposed by the I cells, leading to a locking of I cells to a slightly earlier phase in the gamma cycle than E cells.

Neurophysiological recordings in V1 during stimulus contrast manipulations provide data that permit to increase our understanding of gamma generating mechanisms. A common finding is that stimulus contrast enhancements yield nearly linear increases in gamma frequency in LFP recordings (Jia et al., 2013; Ray & Maunsell, 2010; Roberts et al., 2013) as well as monotonic increases in spike rate (Hadjipapas et al., 2015). The effect of contrast on gamma frequency reflects the fact that contrast is a proxy of excitatory drive (E-drive) (see (Hadjipapas et al., 2015) and references therein). High excitatory drive will overcome decaying inhibition faster, in turn generating gamma at a higher frequency. Thus, contrast affects crucial network interactions in PING models. However, we found gamma power in LFP to saturate or even decrease at high contrast (Hadjipapas et al., 2015; Roberts et al., 2013). This finding has not been widely studied and is poorly understood. Some degree of power decay/saturation was also present in another study in awake monkeys ((Ray & Maunsell, 2010), see figures 1E, 1H)), although this was not commented upon by the authors. Power decay at the highest contrasts has also been observed in anesthetized monkeys by (Jia et al., 2013) in their figure 2C-2D, but without further discussion of that finding. Another, similar study in anesthetized monkeys, however, did not observe power decay (Henrie & Shapley, 2005).

**Figure 1:**
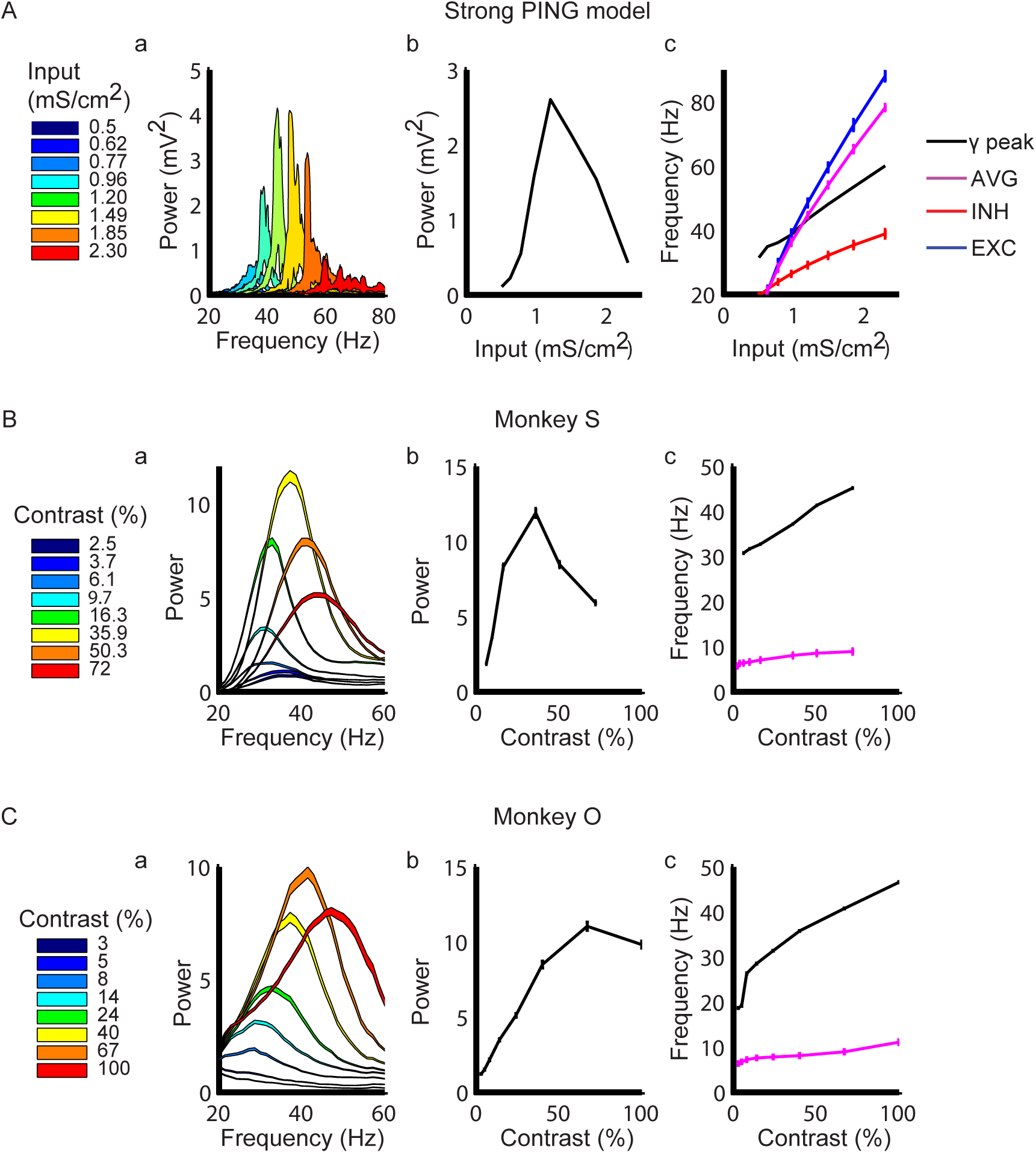
Contrast-dependent Modulation of LFP and spikes in V1 modelled through strong PING and observed in two monkeys (see Roberts et al., 2013). **Aa** Strong PING model. Power spectra of model outputs for different input levels (contrasts, colour code)). A clear frequency shift of spectral gamma response as a function of contrast is observed. **Ab** Power as a function of contrast-strong PING model. From the power spectra in **Aa** two parameters are extracted: peak power and peak (power) frequency. Peak power is plotted as a function of contrast. A nonmonotonic power modulation with contrast is observed. **Ac** Gamma peak frequency (black), average population spiking rate (E&I cells, denoted as AVG) in magenta, as well as E cell (EXC) in blue and I cell (INH) in red spiking rate as a function of input strength (contrast) in a strong PING model. Conventions in **Ba-C** and **Ca-c** as in Aa-c, black line indicates LFP-derived gamma peak frequency and magenta line indicates empirical average single unit rate. **Ba-c** Corresponding observations in monkey S. **Ca-c** Corresponding observations in monkey O. Curve width in Aa, Ba, Ca indicates variability (mean ± SEM). Spectral responses in the monkeys were normalized by baseline. Note that slightly different contrast ranges were used in the two monkeys (see legends). LFP data represent averages over 8 contacts x 17 sessions x average 62 trials in monkey S, and 16 contacts x 45 sessions x average 36 trials in monkey O. Spiking data come from single units isolated from the same datasets (57 in monkey S, and 331 in monkey O).

**Figure 2:**
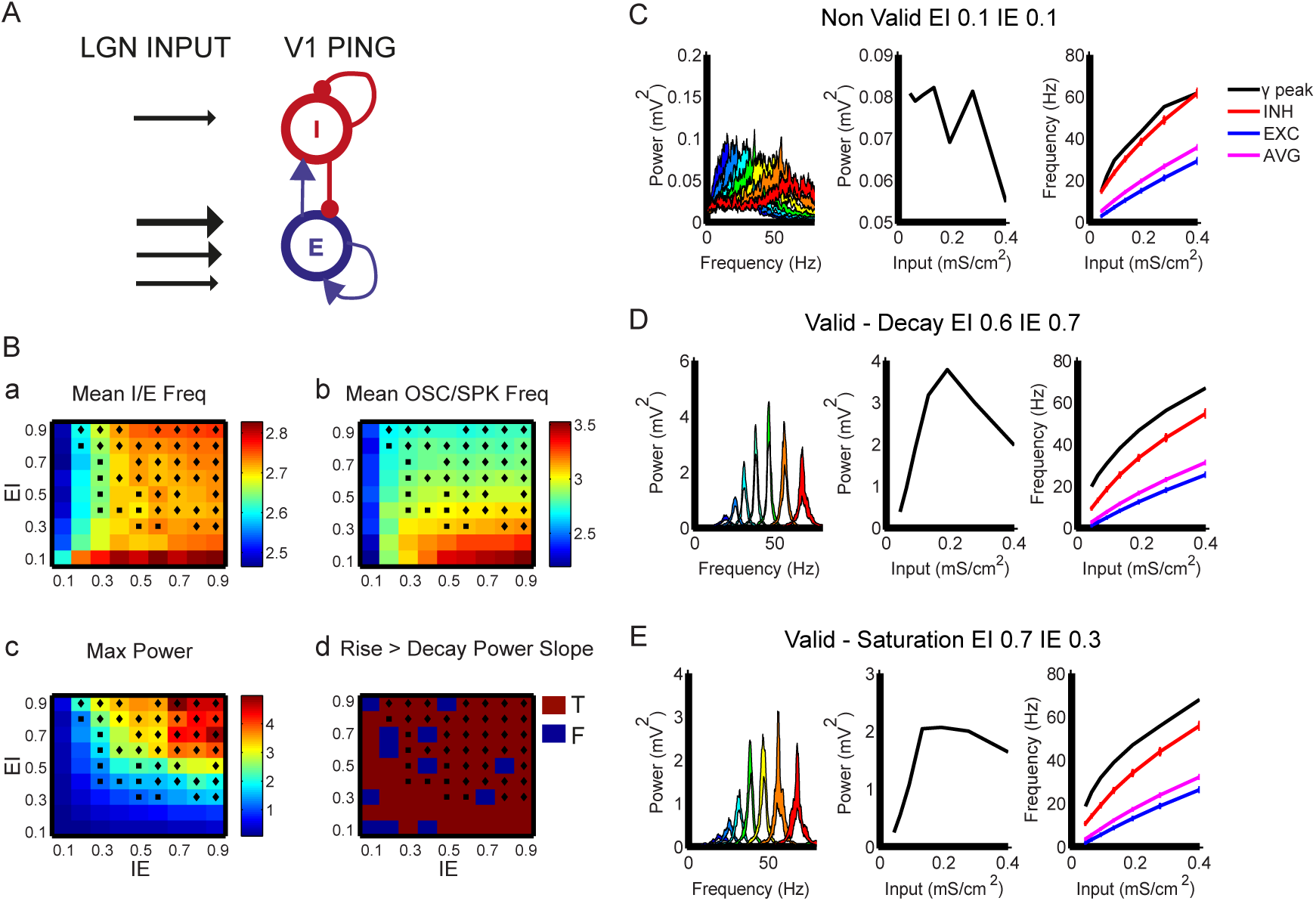
Weak-PING Model Validation. **(A) Network Diagram of PING network.** The model contained regular-spiking E cells (blue) and fast-spiking I cells (red) modelled as Hodgkin-Huxley type single compartment neuron models. Connections within and between populations were set randomly (through voltage-dependent synapses) with a certain probability of connection (EI, IE, EE, II). The LGN afferent input was modelled in terms of Poisson spike trains. The strength of input to the excitatory cells was varied as per conductance magnitude to represent stimulus contrast-dependent input, illustrated by the black arrows of different weight. Input to I cells was of the same magnitude across all contrasts as illustrated by a single black arrow. **(B) Parametric Exploration for E-to-I and I-to-E connection probabilities**. Panels/Colour surfaces correspond to the following criteria (a) #1c, (b) #2, (c) #3b (d) #5 as listed in Table 4, across all possible EI and IE connection probabilities explored here. The horizontal and vertical axes correspond to IE and EI probabilities of connection. Each coordinate (IE, EI) corresponds to a weak PING model with its connection probabilities varied. The output of this model was processed to extract the model output observables that corresponded to the empirical criteria listed above. Valid networks (satisfying simultaneously all criteria) are denoted by a diamond symbol for networks exhibiting power decay and with a square for networks exhibiting power saturation. Colour intensity signifies the value of the criterion in each figure. (**C)-(E). Examples of valid and non-valid weak PING networks**. Same conventions as in figure 1. All results plotted as mean ± SEM. LFP spectral response in three representative networks illustrating the LFP power spectrum for shifting contrasts (left), the peak power as extracted from the spectral response per each contrast (middle) and the peak gamma frequency and average firing rate of all the cells superimposed (right). In the first column power spectra shown in hotter colours correspond to higher contrast stimulation, bluer colours correspond to lower contrast conditions. A non-valid network **(C)** is shown which failed to generate oscillations with high enough power and two valid networks showing a frequency-dependent spectral shift with contrast (left), a non-monotonic power rise and either decay **(D)** or saturation **(E)** with increased contrast (middle).

We should note that beyond the possible effects of anesthesia on power saturation/decay, the stimuli also differed significantly among studies, possibly explaining differences in results. Nevertheless, when considering the two studies using awake monkeys (Ray & Maunsell, 2010; Roberts et al., 2013), power saturation or decay appears to be the predominant observation.

**Table 1:**
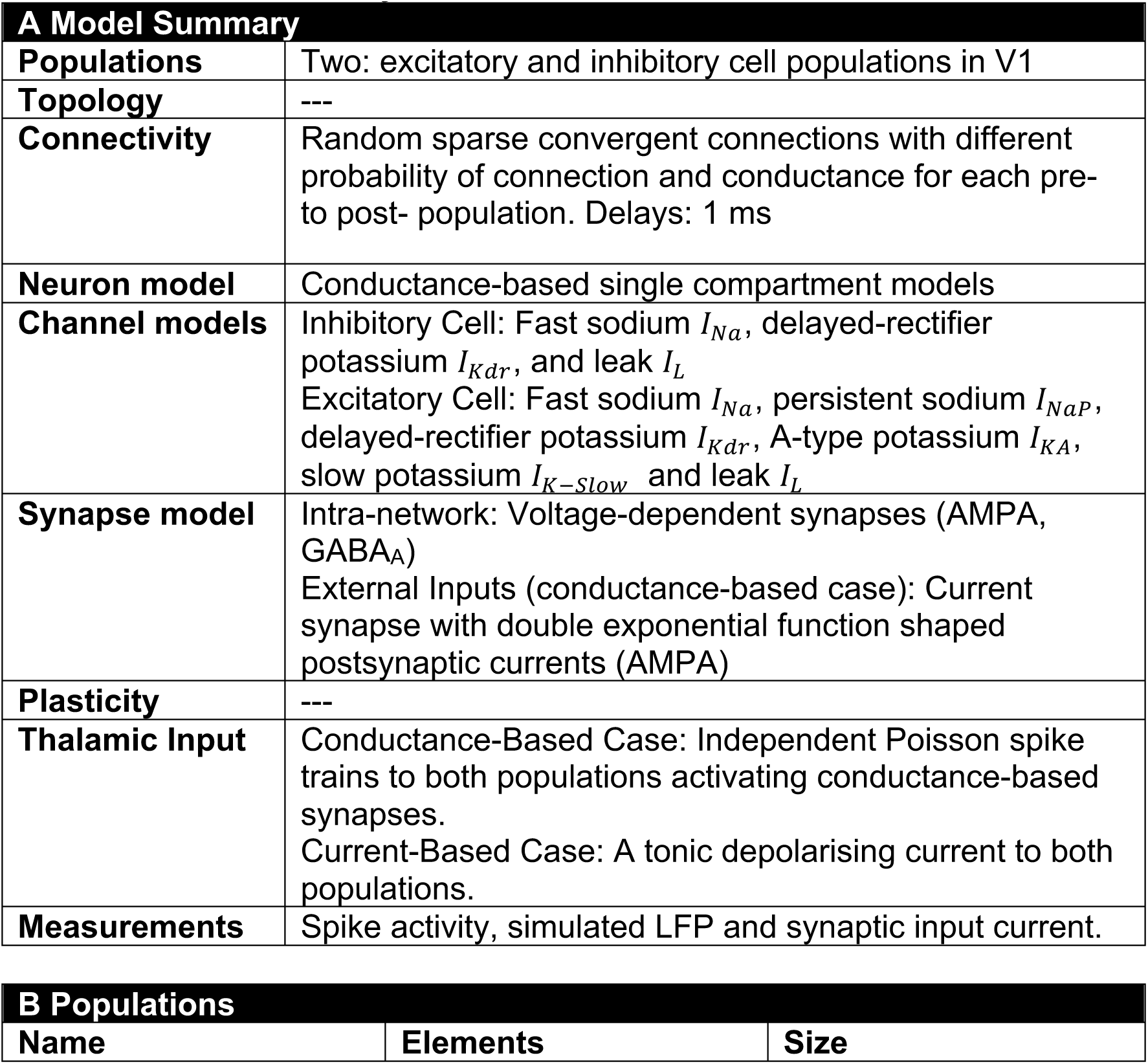
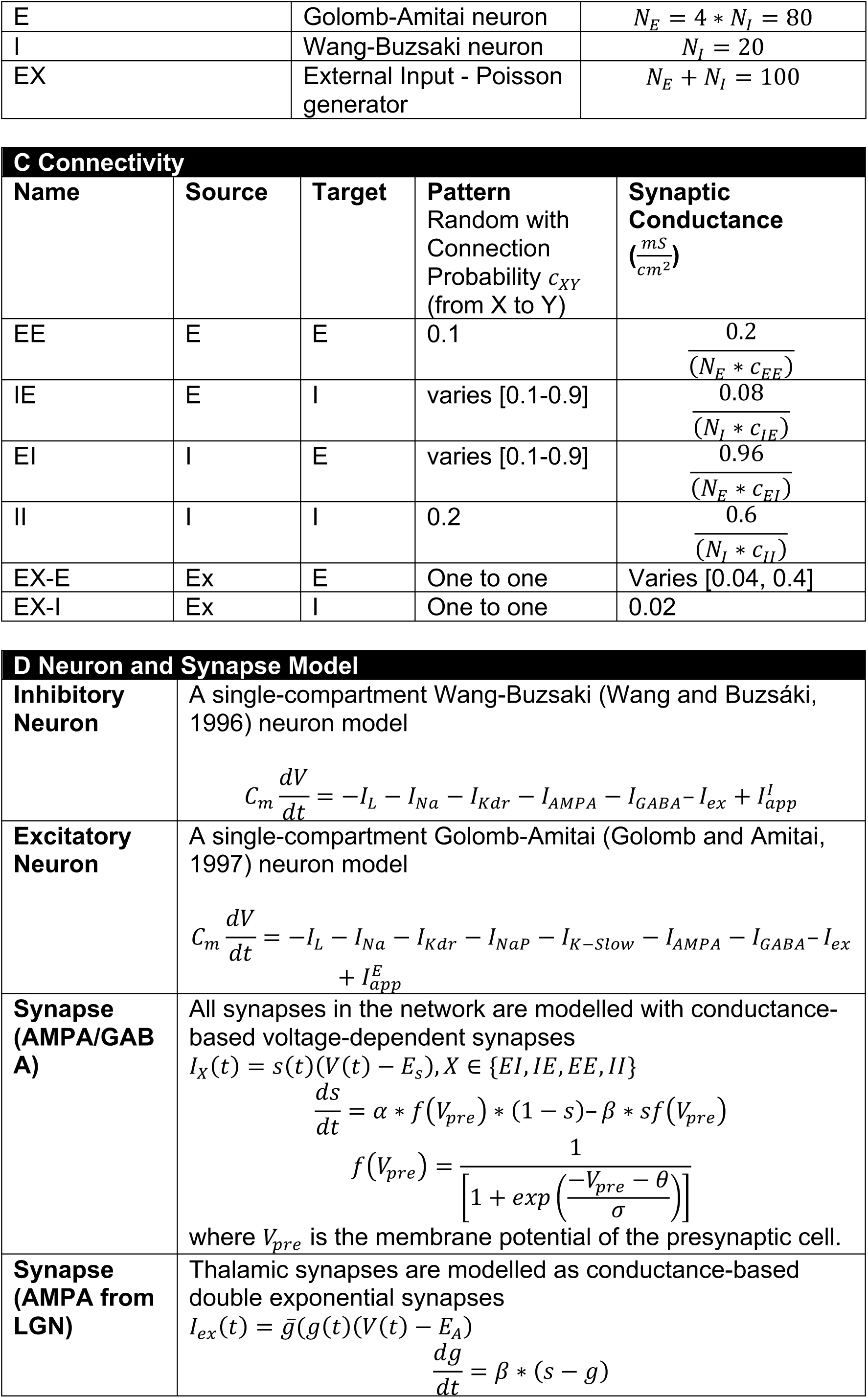
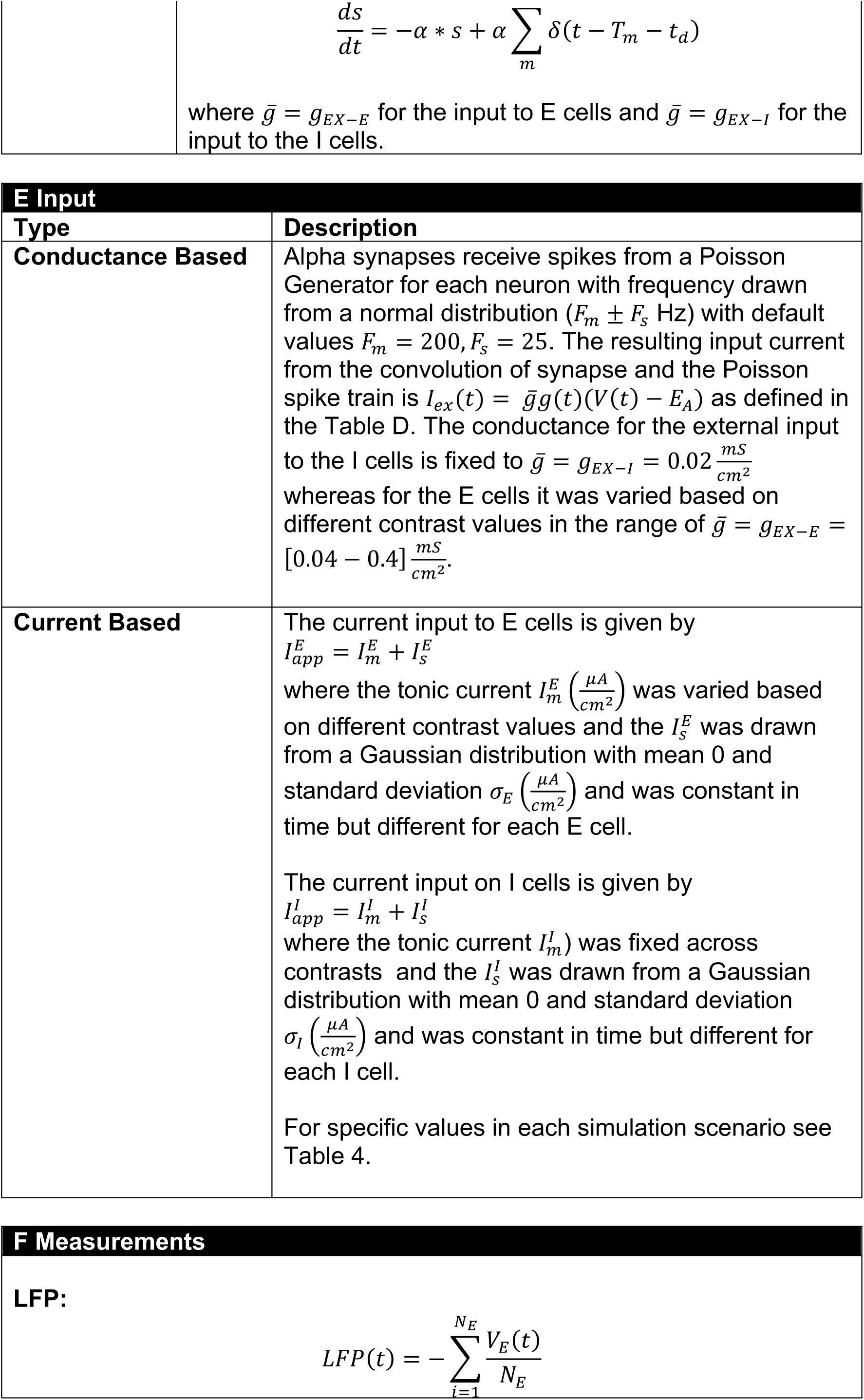
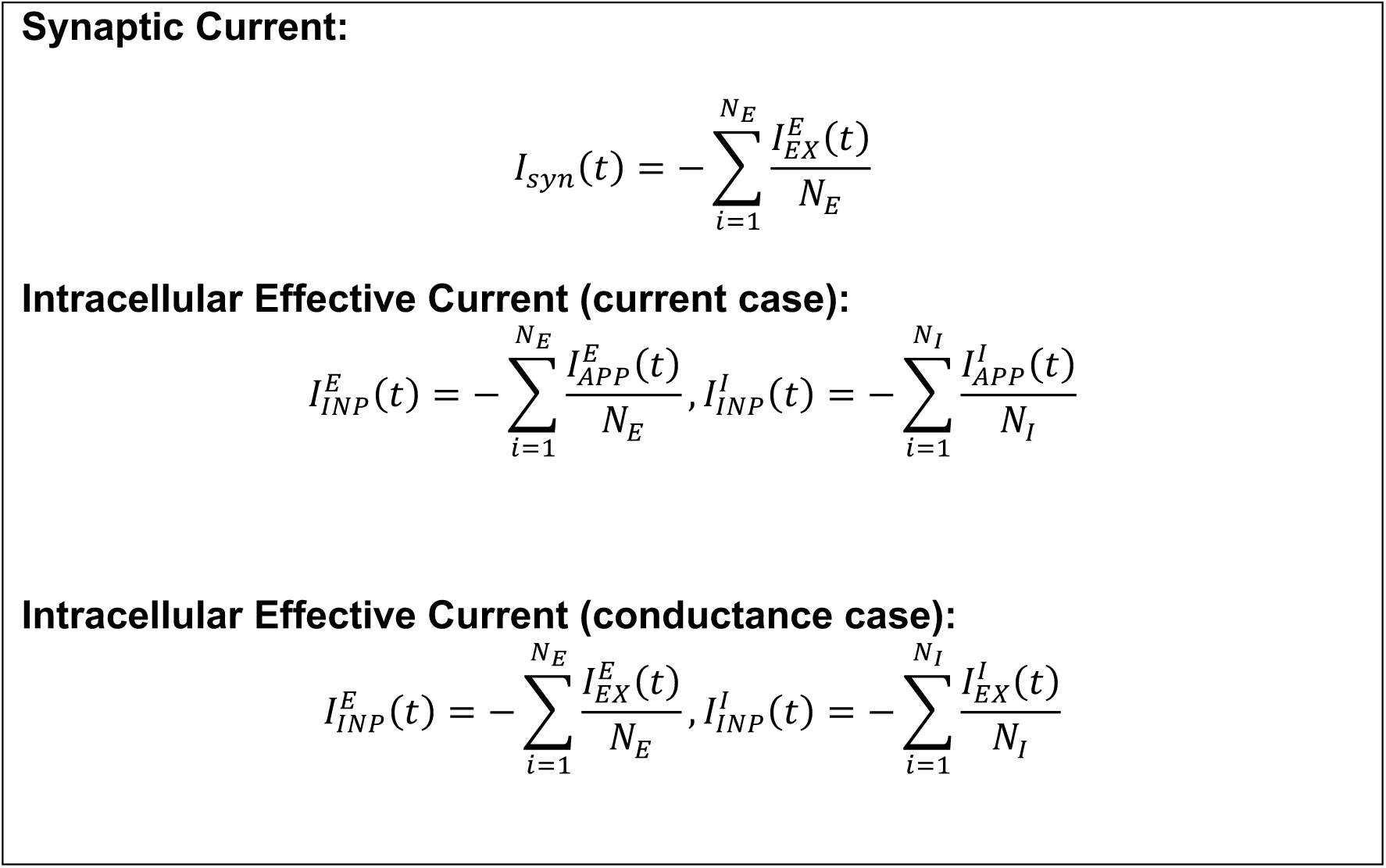
Model Summary.

**Table 2:** Synaptic Parameters.

|  | AMPA | GABA-A | Thalamic AMPA |
| --- | --- | --- | --- |
| $\alpha(ms^{-1})$ | 1.25 | 0.1 | 1.0 |
| $\beta(ms^{-1})$ | 2 | 5 | 5.2 |
| $\sigma(mV)$ | 2 | 2 | - |
| $\theta(mV)$ | -20 | 0 | - |
| $E_{syn}(mV)$ | 0 | -80 | 0 |

**Table 3:**
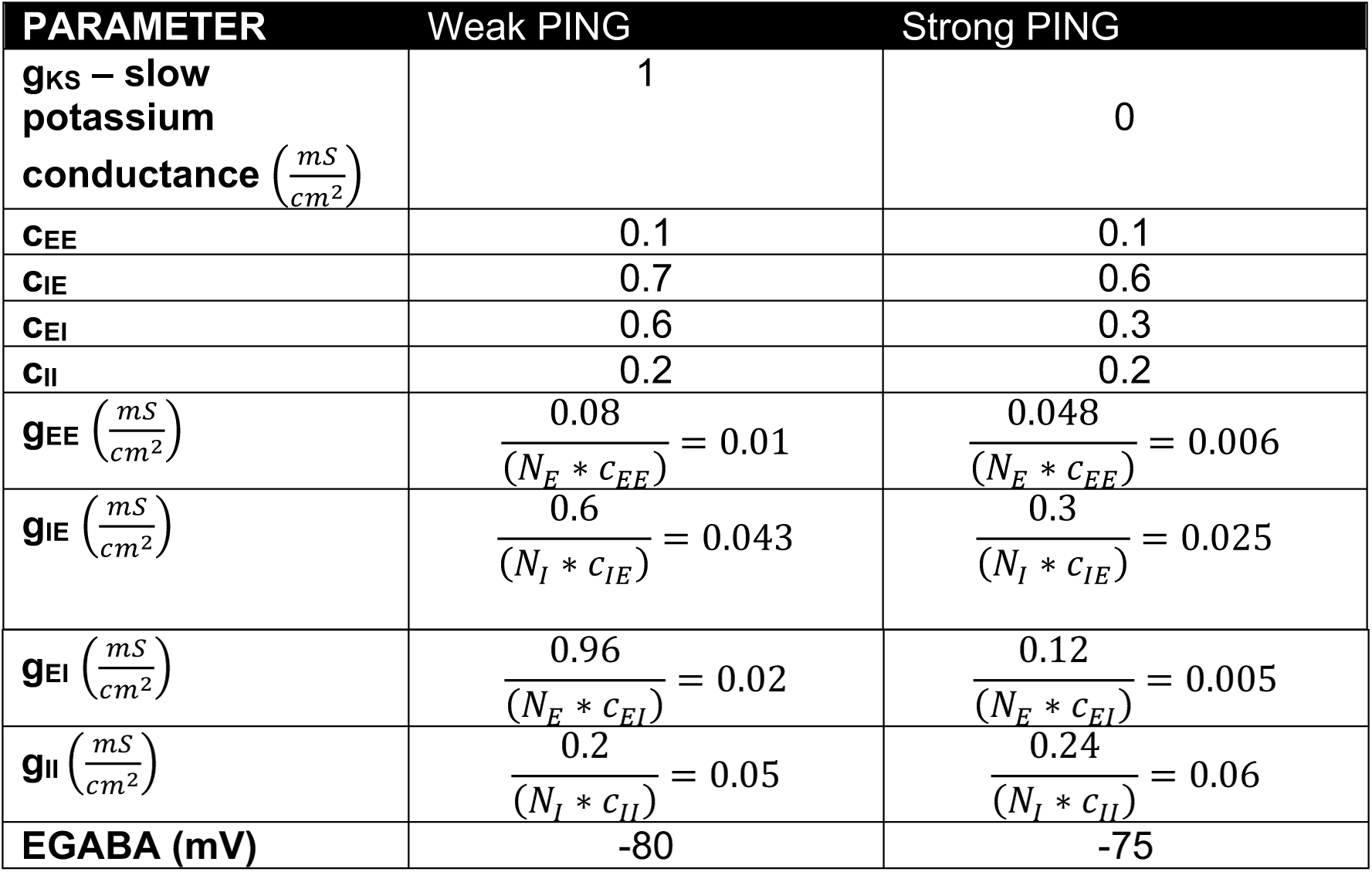
Strong vs Weak Peak with current based input – Main parameters that differ.

**Table 4:** Input Parameters.

| | **Strong PING Current**<br>$\left( \frac{\mu A}{cm^2} \right)$ | **Weak PING Conductance**<br>$\left( \frac{mS}{cm^2} \right)$ | **Weak PING Current**<br>$\left( \frac{\mu A}{cm^2} \right)$ | **Weak PING Current – varying $\sigma_E$**<br>$\left( \frac{\mu A}{cm^2} \right)$ |
| --- | --- | --- | --- | --- |
| **INPUT to E** | $I_m^E$ spaced logarithmically in [0.5,2.3]<br>$\sigma_E = 0.1$ | $g_{EX-E}$ spaced logarithmically in [0.04,0.4] | $I_m^E$ spaced logarithmically in [0.3,4.0]<br>$\sigma_E = 0.01$ | $I_m^E$ spaced logarithmically in [0.3,4.0]<br>$\sigma_E$ spaced logarithmically in [0.01,3.0] |
| **INPUT to I** | $I_m^I = 0.5$<br>$\sigma_I = 0.1$ | $g_{EX-I} = 0.02$ | $I_m^I = 0.3$<br>$\sigma_I = 0.01$ | $I_m^I = 0.3$<br>$\sigma_I = 0.01$ |

PING networks have often been used to model selected aspects of empirical data, at the cost of producing unrealistic network behaviour in other aspects. For example (Roberts et al., 2013) used a strong PING spiking network model that replicated the frequency shift and spectral power saturation/decline that they observed with increased contrast in monkey V1. However, more detailed analysis of this model, reveals unrealistically high firing rates especially for E cells (see results). (Lowet et al., 2015) replicated the same results using a weak PING model with a ring-based topology but applied a long (non-physiological) synaptic time constant for GABA. (Jia et al., 2013) have implemented a population firing rate model that captured the frequency shift but not the power decay as observed in the data. (Mazzoni, Panzeri, Logothetis, & Brunel, 2008) using a PING model captured the behaviour observed in (Henrie & Shapley, 2005) thus not examining the case of power decay/saturation. Hence, existing models have focused on specific phenomena of interest while allowing unrealistic parameters or network behaviour in aspects outside the focus of interest. This shows that even in relatively simple models, many parameters are unknown and are set based on practical considerations or by convention.

To arrive at more realistic PING models, a broader range of parameters needs to be simultaneously constrained by data. To achieve that, we used the extensive data of (Roberts et al., 2013), as well as an existing dataset, not previously reported, which showed power saturation or reduction at high contrasts. Contrary to earlier modelling approaches, we constrained a spatially-undifferentiated PING network through a *combined* exploration of multiple network parameters aimed at approaching a *comprehensive* set of empirical spiking and spectral observations. This was done by taking the following steps:

1. We first characterized the observed gamma LFP spectra in terms of the parametric effects of E-drive (contrast) on the power and frequency of spectral distributions. In addition, we took into account parametric effects of contrast on firing rate, and incorporated estimates of the different firing rates in E and I cells recorded in other studies (Contreras & Palmer, 2003). The sparse relationship between spiking and the LFP gamma oscillation limited the category of plausible models to the weak PING category.
2. For this limited subset of models, we determined combinations of EI and IE connectivity strengths so that manipulations of input parameters in the model replicated the empirical effects of contrast on gamma frequency and power, while also producing realistic neuronal spiking. Within EI/IE parameter space, specific parameter combinations led to either power saturation or power decay at high contrasts replicating experimentally-observed variability between monkeys.
3. Next, using an example of a validated model characterized by strong power decline at high contrasts, we showed that both the mean input level and heterogeneity of input to E cells have to increase with input strength (contrast) in order to generate realistic network behaviour.
4. In further analysis of the requirements of the input to I cells, we discovered that the input to I cells has to be moderate in magnitude and should *not* be modulated by contrast.
5. Investigating a valid model, we then identified possible mechanisms underlying the empirically observed power saturation and decline at high contrast. In the chosen example of a valid model, the power decline was related to oscillation destabilization at high input strength contrasts (E-drive). In this destabilization, both E and I populations played a role, through a mechanism that is distinct from those described in the literature so far.
6. Finally, we used the validated model to investigate discrepancies in the literature concerning the effect of GABA on gamma oscillations such as frequency and power.

We propose that the model derived in our study will be useful for other modelling studies, and that our approach to the empirical constraining of PING models can be expanded when richer empirical datasets become available. As local gamma networks are the building blocks of larger networks that aim to understand complex cognition through their interactions, there is considerable value in improving our models of these building blocks.

## RESULTS

### 1. Weak or strong PING?

We first considered the empirical validity of a strong PING model as used in (Roberts et al., 2013), with the parameter settings listed in **Table 3**. Note that (Roberts et al., 2013) used an LFP proxy based on the firing rate of E cells, whereas in the present work, we used a membrane potential LFP proxy (see Methods). The strong PING model, irrespective of the LFP proxy, reproduced the principal features of the empirical LFP spectral response. Increased contrast shifted LFP frequencies towards higher values (**Figure 1Aa**), whereas after an initial power increase with contrast, a further contrast increase led to power decay (**Figure 1Ab)**. In the light of the similarity of empirical and simulated data, one could conclude that a strong PING model provides a plausible mechanism for the generation of the empirical data.

However, the model unit spiking showed characteristics that were less plausible. E cells (blue line in **Figure 1Ac**) showed much higher firing frequencies than I cells (red line in **Figure 1Ac**). Moreover, E cells (blue line) showed firing rates that were higher than the LFP oscillation frequency (black line), which at high inputs pulled the average spiking rate (magenta line) above the LFP oscillation. This would be highly surprising, and we therefore verified monkey LFP and V1 spiking behaviour as a function of contrast in a dataset used in an earlier paper from our group (Roberts et al., 2013), as well as in an existing dataset, not previously presented.

To evaluate the plausibility of the strong PING model, we attempted to classify single neurons recorded in monkey V1 into excitatory and inhibitory subcategories based on spike shape. This has been successfully done in macaque V4 (Nandy, Nassi, & Reynolds, 2017; Vinck, Womelsdorf, Buffalo, Desimone, & Fries, 2013) and in cat area 17 (Moca et al., 2014). Spike locking behaviour can be used to distinguish between PING and ING. For example, the observed phase-lead of excitatory cells over inhibitory cells in macaque area V4 (Vinck et al., 2013) and cat (Moca et al., 2014) support PING rather than ING. Likewise, cycle-by-cycle strong locking is indicative of strong PING, whereas sparse locking of E cell spiking to the LFP gamma rhythm is indicative of weak PING. Unfortunately, we were unable to come to a satisfactory classification of the V1 spiking cells in our data (see Methods). Nevertheless, the population spiking rate proved to be a valuable tool to constrain PING models.

**Figures 1B and 1C** show for two monkeys the LFP spectral and spiking response to a range of grating contrasts. In the two monkeys, gamma spectra were contained within a 20-80 Hz frequency range and spectra at low contrast were characterized by very low power. The spectral response was furthermore characterized by a saturation or decay in power in the high contrast range (**Figure 1Bb** and **1Cb**, and a linear increase in LFP gamma frequency (at peak power) as a function of contrast (black line **Figure 1Bc, 1Cc**). Population spiking rate showed a modest, linear increase with contrast (magenta line **Figure 1Bc, 1Cc**).

A highly relevant observation in both monkeys was that the population spiking rate was very low compared to LFP gamma frequency (compare magenta and black lines in **Figures 1Bc, 1Cc**). This observation disqualifies the strong PING model used in (Roberts et al., 2013). Instead, the data suggest a sparse rhythm in the neural population in which only a subset of neurons participate in the population oscillation in a given cycle. This idea is in line with the notion of sparse rhythms and, thus, with a weak-PING model as the underlying gamma generation mechanism (Börgers, Epstein, & Kopell, 2005; Wang, 2010). The sparseness of single unit firing with respect to the population oscillation can be characterised by the ratio between LFP gamma frequency and single unit firing rate. This was also extracted from the empirical data. To inform the relative firing rates of E and I neurons as a function of contrast we utilised work by (Contreras & Palmer, 2003), from where we extracted the relative ratio of I/E firing rates.

When attempting to build a PING type model, input and connectivity parameters are of utmost importance for model behaviour, yet they remain unobserved in the physiological counterpart of the model due to lack of direct and specific empirical data to guide as to their choice. Nonetheless, an alternative approach exists, whereby such parameters can be constrained by empirical data by maximizing the match between model output and observed behaviour in the empirical data. This entails using a set of criteria derived from the empirical data that model output must meet. Thus, in our case, the requirement is that model output must capture the most salient contrast-related modulations of gamma oscillations in V1 both at the LFP and single unit spiking level. To this end we derive a set of 5 *criteria* for model validation (as illustrated in **Figure S1** and described in detail in Methods - Table 5):

**Table 5:** Empirically-obtained Validation Criteria for models.

| **Criteria** | **Description** |
| --- | --- |
| **#1:** I cells fire more frequently than E cells. | We fit the firing rates linearly across contrasts ($Inp$) of the E and I cell populations ($FE$ and $FI$) with<br>$FE_{fit} = s_E * Inp + a_E ,$ $FI_{fit} = s_I * Inp + a_I ,$<br>and consider the following conditions:<br>The intercept of the $FE_{fit}$ must be less than the intercept of the fit to $FI_{fit}$, ($a_E < a_I$).<br>The slope of $FI_{fit}$ must be bigger than the slope of $FE_{fit}$ ($s_I > s_E$)<br>The average of the I cells (actual) firing rate must be at least 2.5 times higher than the average E cell firing rate ($\langle FI \rangle \geq 2.5 * \langle FE \rangle$). |
|  | All conditions a-c must apply for criterion #1 to be deemed valid. |
| **#2:** LFP oscillation frequency is higher than the population firing rate | The average LFP oscillation frequency ($FO$) must be at least 2.3 times higher and less than 6.3 lower than the average firing rate of the whole population ($FP$), $2.3 * \langle FP \rangle \leq \langle FO \rangle \leq 6.3 * \langle FP \rangle$. |
| **#3:** LFP first and peak power | (a) The peak power must be larger or equal to 1.0.<br>(b) The power at the lowest contrast must be less than 0.5. |
| **#4:** LFP oscillations in gamma range. | The LFP oscillations must be in the gamma range (15-80 Hz). |
| **#5:** LFP peak power rise is faster than its decay. | We linearly fit the slope for the power rise (POR) and the decay (POD) of the power with<br>$POR_{fit} = s_R * FOR + a_R,$ $POD_{fit} = s_D * FOD + a_D.$ We then checked that the rise slope was steeper than the decay slope $s_R \geq s_D$. |

1. I cells should fire at least 2.5 times more frequently than E cells while both should increase their output linearly within the contrast range used. The data from (Contreras & Palmer, 2003), suggested that the ratio of I to E cell firing rates could be as large as fivefold. However, from that data no minimum condition could be extracted. In extensive simulations that we ran the minimum ratio was close to 2.5, hence we set this parameter at this value.
2. The LFP gamma peak frequency must fall between 2.3 and 6.3 times the average single-unit firing rate, in line with empirical observations. This was based on the mean of the two monkeys +/- 3 standard deviations.
3. The power at the lowest contrast should be smaller than a threshold value (0.5) and the maximum power should be greater than a threshold value (1.0); These two limits were empirically applied to capture the qualitative observation that at low contrasts gamma power is low, and to ensure that a gamma oscillation indeed exists in the model outputs respectively. As the scales of the model and data were different and thus data could not be readily utilised for these limits, these values were chosen based on several model simulations such that they best captured the qualitative behaviour observed in the data.
4. In the input to power-function, the flank to the left of the peak (ascending slope) should be less steep than the right flank (descending slope), based on the mean function of the two monkeys.
5. LFP oscillations should fall within the gamma range (15-80 Hz)., also based on the mean of the two monkeys. The gamma range is often considered to start above 20 Hz, however, for low contrasts in our experiments we observed oscillations in the 15-20Hz.

Note that this approach does not represent an attempt to accurately fit the available empirical data but rather to qualitatively capture the most salient features of the modulation of gamma oscillation by contrast.

### 2. Exploring Empirically Constrained Weak PING Models

We constructed a spatially undifferentiated weak PING model, according to the 5 criteria derived in the previous section. This PING model can be considered as a much simplified local network in V1. The network consisted of 100 Hodgkin-Huxley type cells; 80 excitatory cells and 20 inhibitory cells. These were randomly connected, whereby all possible connections (E to I (EI), I to E (IE), I to I(II), E to E (EE)) occurred with a certain probability (**Figure 2A**). Synapses were modelled by a voltage-dependent model, representing fast excitatory and inhibitory currents with realistic time courses. Both populations received a noisy external excitatory input reflecting the LGN afferent input. To obtain weak PING the E cell frequency input current (f-I) curve and the synaptic conductances were adjusted to facilitate sparse firing of E cells (see Methods for details).

The crucial connectivity parameters to achieve oscillations in PING models are the EI and IE connection probabilities (Buzsáki & Wang, 2012; Lee & Jones, 2013). We applied a 2-dimensional parameter manipulation, in which the EI and IE connection probabilities were manipulated systematically. The resulting model outputs were evaluated against our 5 criteria in order to select valid connectivity parameters. In **Figure 2B**, each coordinate (IE, EI) corresponds to a different set of connection probabilities. The output of the simulated models for each coordinate was processed to extract the model output observables that corresponded to the empirical criteria. The values of these model observables are color-coded in the surface plots in **Figure 2Ba-d**. Only networks that simultaneously satisfied *all* criteria were considered empirically valid and are indicated with a diamond (networks exhibiting power decay) or square symbol (networks exhibiting power saturation) in the panels in **Figure 2B**. These panels correspond to the following criteria (see Methods for quantitative details): Criterion **#1:** I cells fire more frequently than E cells (**Figure 2Ba**); Criterion **#2:** the LFP oscillation frequency is higher than the population firing rate (**Figure 2Bb**); Criterion **#3:** Maximum power is greater than a threshold value (**Figure 2Bc**); Criterion **#4:** LFP peak power rise is faster than its decay (**Figure 2Bd**.). Note that Criterion **#5**, stipulating that oscillations must be in the gamma range (15-80 Hz), was satisfied for all but one set of connection probabilities ((EI 0.1, IE 0.1), and is not illustrated in **Figure 2B**. Three cases of networks with different behaviour are illustrated in **Figure 2C-E**. The first is a non-valid network (EI 0.1, IE 0.1) with very low overall power (panel C), the second a valid network (EI 0.6, IE 0.7) exhibiting fast decay (panel D), and thirdly a valid network (EI 0.7, IE 0.3) exhibiting saturation (panel E). In all three models, and in line with weak PING, neurons fired sparsely, I cells exhibited a higher firing rate than E cells, and the LFP oscillation frequency was higher than the average firing rate of the neurons. The network shown in **Figure 2C** was invalid due to the weak EI and IE connectivity. Note that model parameters including (but not limited to) EI and IE connectivities could be set to simulate network outputs not showing power saturation or decay (data not shown), but here we focused on constructing valid simulations for the empirical data in our own hands.

Going beyond these three examples, we report the following general observations: All tested network parameter sets for EI and IE connection probabilities for networks were valid with respect to Criterion #1 (I cells fire more frequently than E cells) and #2 (LFP oscillation frequency is higher than the population spiking rate). However, changes in EI and IE connection probabilities distinguished valid from invalid models based on the other 3 criteria. Notably, valid models were not a few, but rather many, which within a broad range of EI and IE connection probabilities reproduced power decay (diamonds in **Figure 2B**) or power saturation (squares in **Figure 2B**) at high contrast. In addition, the models exhibiting saturation and decay occupied separate regions in the EI/IE parameter space. Specifically, when the sum of EI and IE connection probabilities was greater than 1, decay was observed; when this sum was less than 1, saturation was observed. When this sum was equal to 1 both cases were observed for different EI and IE parameter combinations. In summary, the described validation process indicated the importance of a specific relative ratio between EI and IE connection probabilities for the network to exhibit realistic (empirically valid) behaviour. In the following sections we used a specific valid instantiation (in terms of connection probability parameters) defined as the *default* weak PING for further investigations (see Table 3 for parameter settings).

### 3. Input Requirements: To obtain realistic behaviour both the mean input strength and heterogeneity across E cells must be modulated by contrast

In the in-vivo organism there is a complex afferent system whereby V1 superficial layers (where gamma is the strongest) receive input across multiple stages (the physical luminance input, the retina, the LGN and the input layers of V1). By the term input we refer here to the outcome of this afferent processing system which reaches the superficial layers in V1.

In the strong PING model simulated in Section 1 (Roberts et al., 2013), the input was current-based (see Methods). However, in our validated weak-PING model, we used conductance-based input. This raised the question of how the nature of input may influence model behaviour. To investigate this, we exposed a strong and a weak PING model (see Table 3 for full set of parameters) to the same current input. Conversely, we took the same weak PING model and compared its behaviour when exposed to current or conductance input. In contrast to E cells, I cells in both models were not subjected to contrast-dependent modulations; they received a low-level, constant drive (see Section 4).

To directly compare the properties of the input in the different cases, we “recorded” the intracellular effective current *I*_*INP*_(*t*) resulting from external (to the model network) input to each cell. For the current-based input case, *I*_*INP*_(*t*) is a tonic current applied to each cell, and for the conductance-based input case, *I*_*INP*_(*t*) is obtained from the convolution of the synaptic model with a Poisson spike train (see Table 1E for more details). We then extracted two descriptive characteristics of the external contrast-modulated input to the entire population. Specifically, the input *magnitude* 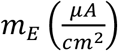 was calculated by taking the average *I*_*INP*_(*t*) across E cells over the duration of stimulation, followed by averaging over trials. Input *heterogeneity* 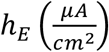 was calculated by taking the standard deviation of *I*_*INP*_(*t*) across E cells per trial for the duration of stimulation, followed by averaging over trials.

In **Figure 3**, rows **A-D** show the model outputs of input manipulations for the selected valid default weak PING model and for the strong PING model of Roberts et al. (2013). Column **a** represents peak power as a function of input. Column **b** represent the spike rates of E, I and all (AVG) neurons and the LFP peak frequency as a function of input. Column **c** shows the input *magnitude m*_*E*_ to E cells as a function of contrast modulation. Column **d** represents the input *heterogeneity h*_*E*_ for E cells as a function of contrast modulation.

**Figure 3:**
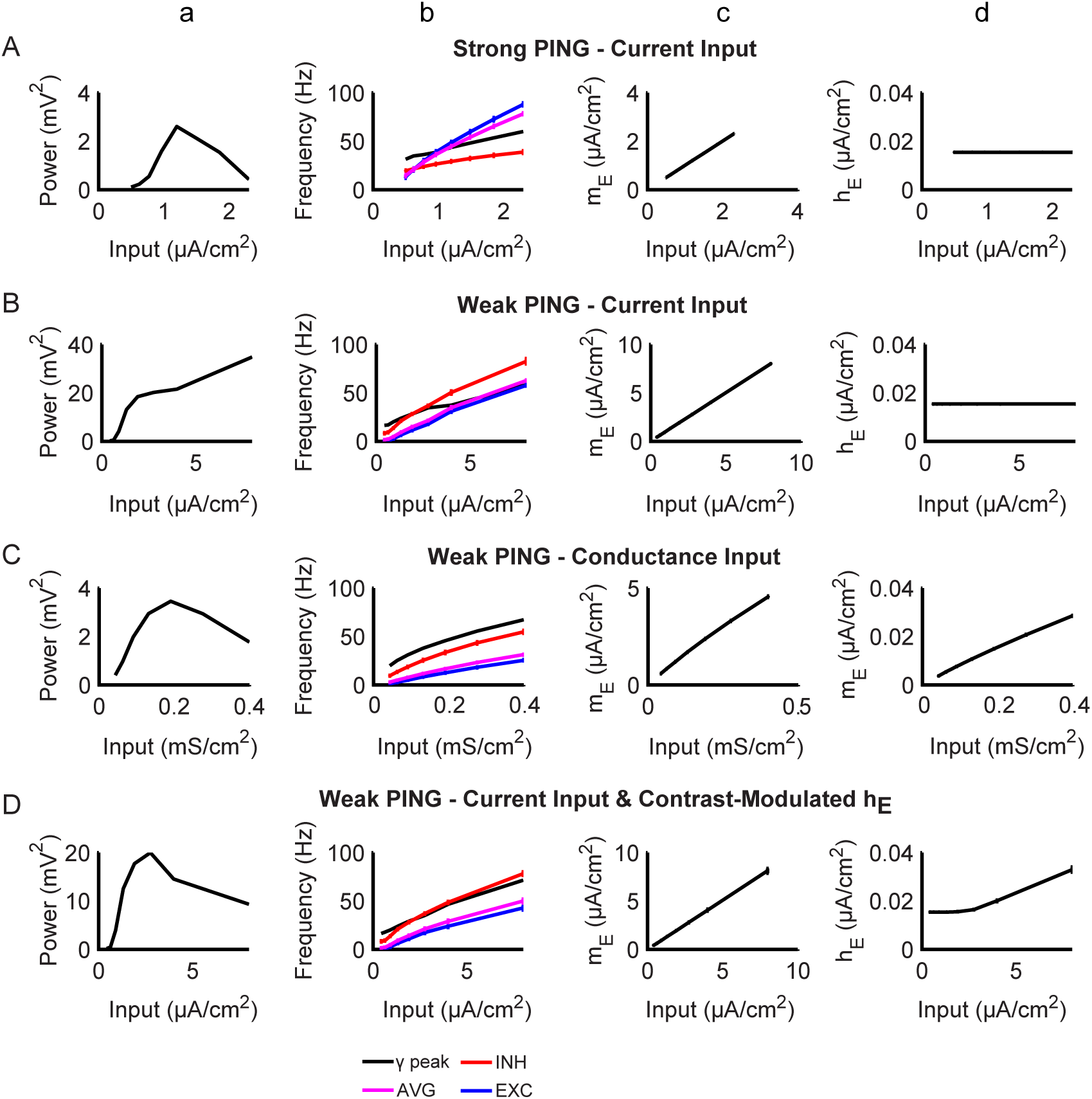
Effects of input properties on E cells. Panels A-D: All results plotted in as mean ± SEM. The first two vertical columns (a and b) show parametric variation of network oscillation features as a function of input strength. The **(a)** column shows gamma peak power of modelled LFP and the **(b)** column the average single unit firing rates for E, I and all cells (AVG) as well as LFP gamma peak frequency (gamma peak) (conventions as in **Figure 1**). The last two columns show properties of the input as modulated by contrast, column **(c)** shows the magnitude (m_E_) and column (d) the heterogeneity (h_E_) of the intracellular effective input current across E cells. Rows represent simulation results from different model types. **Row A** shows **strong PING with current input** (used in (Roberts et al., 2013)), which did not satisfy all criteria as outlined in section 2, although it exhibited the key LFP-based features such as frequency shift and power decay. **Row B** shows a **weak PING model with current input** (current based, mean input (m_E_) varies with contrast). The network failed to exhibit a power decay/saturation at high inputs. **Row C** shows results from a **weak PING model receiving a Poisson conductance-based input** (both mean (m_E_) and heterogeneity (h_E_) of input are modulated by contrast). **Row D** shows a **weak PING model with current input, modified** such that not only the mean (m_E_) but also the heterogeneity (h_E_) of the input across neurons varied with contrast. This resulted in a valid network with similar decoupling mechanism as in the case of Poisson-conductance-based input (**Row C**). Results show that to obtain the power decay/saturation at high contrasts, both the mean (m_E_) and the heterogeneity (h_E_) of input across E cells have to be modulated by contrast.

For the strong PING model with current input (row **A**), only the input magnitude *(m*_*E*_) was scaled with input strength (**Figure 3Ac**), whereas the heterogeneity *(h*_*E*_) was constant (**Figure 3Ad**). This model produced plausible LFP spectral output in the gamma range (**Figure 3Aa**), but at the cost of unrealistic model unit spiking rates (**Figure 3Ab**). Remarkably, as shown in row **B**, we found that the current input with the same characteristics (i.e. contrast-modulated input magnitude and constant heterogeneity) given to the weak PING model led to a power increase without saturation, thus not matching the empirical data (**Figure 3Ba**). Hence, network responses to the same kind of input differed strongly between weak and strong PING models.

We then examined which input properties would restore valid network behaviour for the weak PING model. To this aim, we first returned to the conductance-based input that we had used in Section 2. Note that in the conductance-based model (row **C**), input strength is implemented as conductance strength *g*_*EX*−*E*_ to E cells. For this type of input, an increase in input magnitude will inherently lead to an increase in input heterogeneity (**Figure 3Cc-d**). This is due to the convolution of the conductance *g*_*EX*−*E*_ with the poissonian synaptic input reflected in *g*(*t*) in the synapse model (AMPA from LGN, Table D). The increased input heterogeneity due to the increased input magnitude in the conductance-based case appeared crucial to achieve a realistic level of power saturation, as well as realistic spiking behaviour (**Figure 3Ca-b**). Next, we tested whether contrast-modulated input heterogeneity was crucial for plausible weak PING network behaviour also for the current-based input case (Row **D**). We found that in the current-based input case, the concurrent modulation of input heterogeneity and input strength indeed recovered the power decay behaviour (**Figure 3Da**), while keeping realistic spiking behaviour (**Figure 3Db**).

Thus, an interplay between contrast-modulated input magnitude and input heterogeneity is required for realistic network behaviour. To achieve a better understanding of this interplay we systematically varied parameters governing input magnitude and heterogeneity in weak PING models using either current-based (**Figure 4A**) or conductance-based input (**Figure 4B)**. Peak frequencies tended to increase as a function of input strength irrespective of input heterogeneity for both current-based input (**Figure 4Ab**) and conductance-based input (**Figure 4Bb**). However, peak power showed a very different landscape for current-based input (**Figure 4Aa**) and conductance-based input (**Figure 4Ba**). For the current-based input, the peak power landscape was such that constant heterogeneity at any level would lead to invalid network behaviour, namely, to monotonically increasing peak power as a function of input strength (dashed coloured lines in **Figure 4Ac**). Only when heterogeneity co-varied with input strength (arrow in the power landscape in **Figure 4Aa**) did the model produce valid behaviour of the peak power as a function of input strength (black solid line in **Figure 4Ac**). Whether input heterogeneity co-varied with input strength or not, the model always produced an increase in gamma frequency as a function of input strength (**Figure 4Ad**). **Figure 4Ae** shows that the E-drive given to individual model units is perfectly correlated with the effective average current, measured at the level of E units in the model. **Figure 4Af** shows the input heterogeneity of the effective average current on E cells (dashed coloured lines) over input strength, whereby dashed coloured lines correspond to different but constant levels of input standard deviation σ_*E*_, (the parameter governing heterogeneity, see legend). The same figure also shows the condition in which input strength and heterogeneity co-varied (black line, see also **Figure 3Dd**). For the conductance-based input (**Figure 4B**), there is an inherent positive linear relationship between conductance-based input strength, and heterogeneity of activity in model E cells, which reflects different levels of frequency variability over time in the Poisson spike train (**Figure 4Bf**). This led to the specific power landscape in **Figure 4Ba. Figure 4Bc** shows plausible peak power variations as a function of input strength at all levels of input heterogeneity (coloured solid lines), with as the only exceptions the cases with very high input heterogeneity (dashed orange and red lines). As opposed to peak power variations, peak frequency variations did not differ between different input heterogeneities (**Figure 4Bd**).

**Figure 4:**
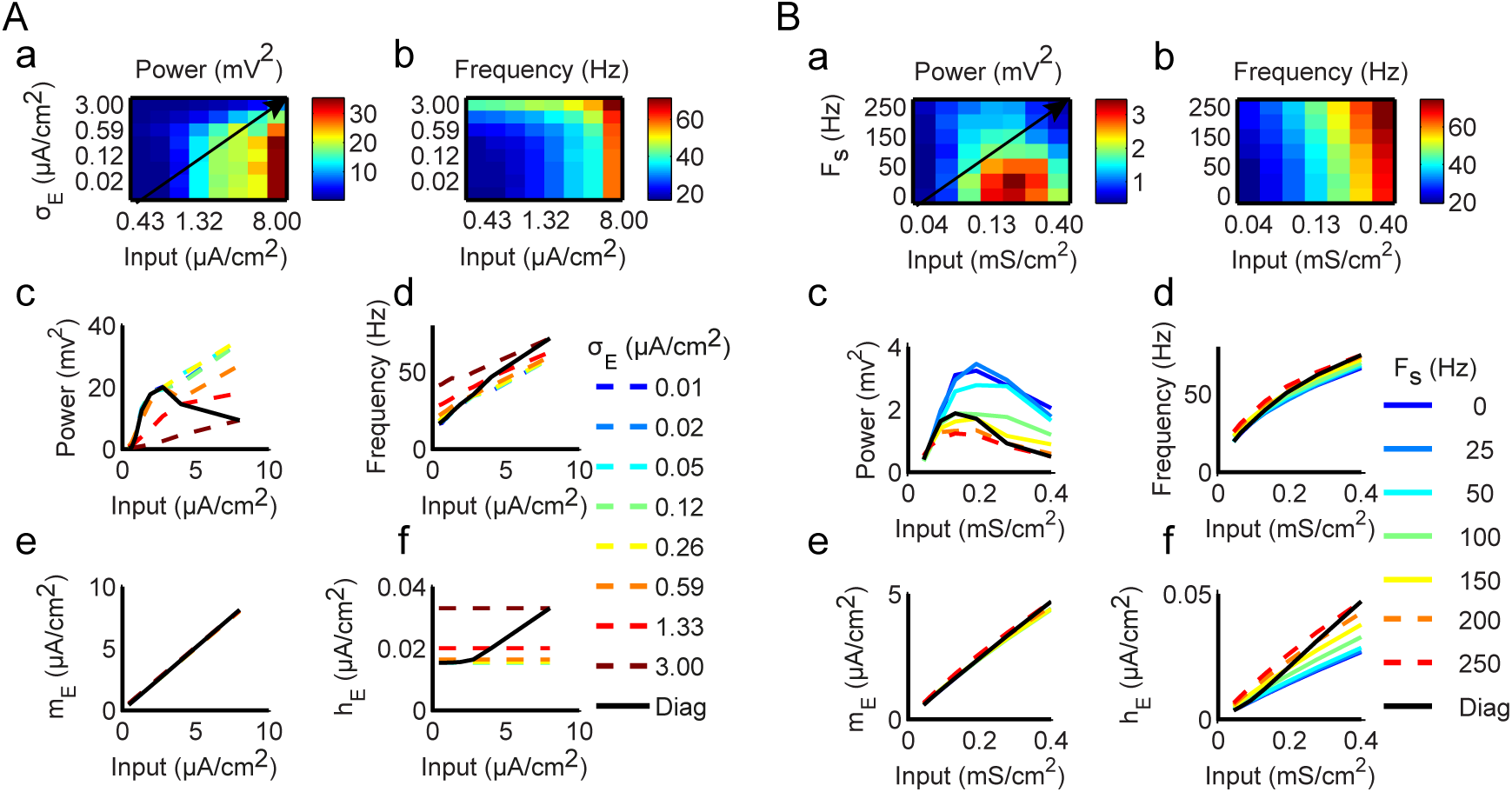
Interplay of Input Heterogeneity and Input Magnitude. **A. Weak PING with current based Input. A(a)** LFP peak power landscape plotted as a function of input strength 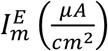, and input standard deviation 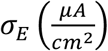, i.e, the parameter that governs the input heterogeneity. Peak power is colour-coded such that highest power is in red. **A(b)** Landscape of gamma peak frequency as a function of input strength 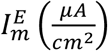 and input standard deviation 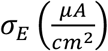. Peak frequency is colour coded from high (red) to low (blue). **A(c)** LFP peak power obtained as a function of input strength for different constant levels of input standard deviation σ_E_. Coloured lines indicate peak power from low σ_E_ values (blue) to high σ_E_values (red). The power spectra correspond to horizontal cross sections through the power landscape in A(a). **A(d)** Peak frequency plotted as a function of input strength, for different conditions in which a constant level of input standard deviation σ_E_ is maintained across levels of input (colour coded from low σ_E_values (blue) to high σ_E_ values (red)). **A (e and f)** Average mean (indicating magnitude - m_E_) (e) and average standard deviation (indicating heterogeneity h_E_) (f) of the intracellular effective contrast-modulated input current across E cells as a function of input strength 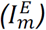. Coloured dashed lines show conditions with different input standard deviations which were however, maintained constant across the input strength manipulation. Note that in all cases where input heterogeneity is constant across input strengths, there is no realistic relation between peak power and input strength, as indicated by dashed lines in A (c-f). Only when σ_E_ increases linearly with input strength (black line in A(f)) also corresponding to black arrow in A(a)), does a realistic power saturation/decay occur (for further details, see text). **B. Weak PING with conductance-based Input. B (a)**. The conventions are the same as in panel A with the difference that the LFP peak power is landscape plotted as a function of parameter 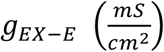 governing input strength and the frequency variability of the poissonian train F_s_(Hz), whereby F_s_ governs the rate of heterogeneity change with contrast. In Panels **B c-f** continuous lines indicate valid, while dashed lines indicate invalid networks.

Overall, our findings indicate that input strength and input heterogeneity in activity levels across E cells are two factors with dissociable contributions to the spectral outputs of the weak PING model. On the one hand, the manipulation of input strength to E cells alone was sufficient to obtain the experimentally observed LFP frequency shift (and the initial increase of power with contrast at low contrasts). On the other hand, plausible spectral power decays/saturations at high contrasts relied on increasing heterogeneity of the input across E cells as a function of input strength. This held true irrespective of whether the input was conductance–based or current-based.

### 4. Input Requirements: To obtain realistic behaviour the input to I cells should be moderate in magnitude and should not be modulated by contrast

We have assumed, as others (Lowet et al., 2015; Mazzoni, Brunel, Cavallari, Logothetis, & Panzeri, 2011; Roberts et al., 2013), that contrast-dependent input from LGN modulates input to the E-cells but not the I-cells, which instead receive the same level of input across all contrasts. However, to our knowledge, this assumption has not been tested, neither in neurophysiological or optogenetics studies, nor in modelling studies. Here, we used the default validated weak PING model with conductance input (**Figure 2D**) to verify the levels of input to E and I cells that so far were set by convention.

### Magnitude of Fixed Input to Inhibitory Cells

We first investigated the role of the magnitude of input to the I cells, which in previous sections was *set at a fixed level* across all contrasts, while the input to E cells varied with contrast. We repeated the simulations for different scenarios each corresponding to a different level of input *g*_*EX*−*I*_ to the I cells (fixed across all contrasts). **Figure 5A** illustrates two cases, one with weak constant input to I cells (left, thin arrow), and the other with stronger constant input to I cells (right, fat arrow). The contrast-modulated input to E cells applied in each case is indicated by the arrows of varied thickness. The magnitude of the input was varied by modulating the conductance (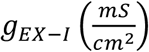 see Methods). Note that because of the nature of conductance-based input, *g*_*EX* − *I*_ increases to I cells will increase both the magnitude and the heterogeneity of the effective input to I cells (as discussed previously for the case of E cells). We plotted LFP peak power (**Figure 5B)** and peak frequency **(Figure 5C)** as a function of input strength for cases of different constant input levels to I cells (colour coded in **Figures 5B-C**). Continuous lines represent valid networks based on the criteria outlined in section 2, while dashed lines correspond to invalid networks. We only obtained valid networks for intermediate values of conductance to I cells 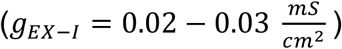 (shown in *continuous* lines). Increased conductance to I cells 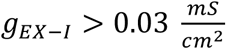 shifted the oscillation frequency into a lower range (<15 Hz) for low contrast values. This also resulted in an overall decrease in power, rendering the network invalid in these case conditions (dashed yellow and orange lines, **Figure 5B**). Conversely, decreased conductance in I cells (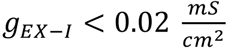 resulted in overall higher power (**Figure 5B**). In this case, we observed an unrealistically high power for the lowest contrast (failing criterion #3b) (blue dashed lines **Figure 5B**) and an unrealistic average firing rate ratio of the two populations (failing criterion #1, not shown). Notably, because of the conductance-based input to E cells, input magnitude to the I cells will increase concomitantly with variability among the I cells as a function of increasing contrast within the higher contrast range. In turn, I–cells can be expected to increasingly desynchronize at high contrasts, which in turn must have led to E cells producing gamma oscillations at decreased power. This suggests that I cells have a key pacemaker role but their synchronising potential weakens at high input strength.

**Figure 5:**
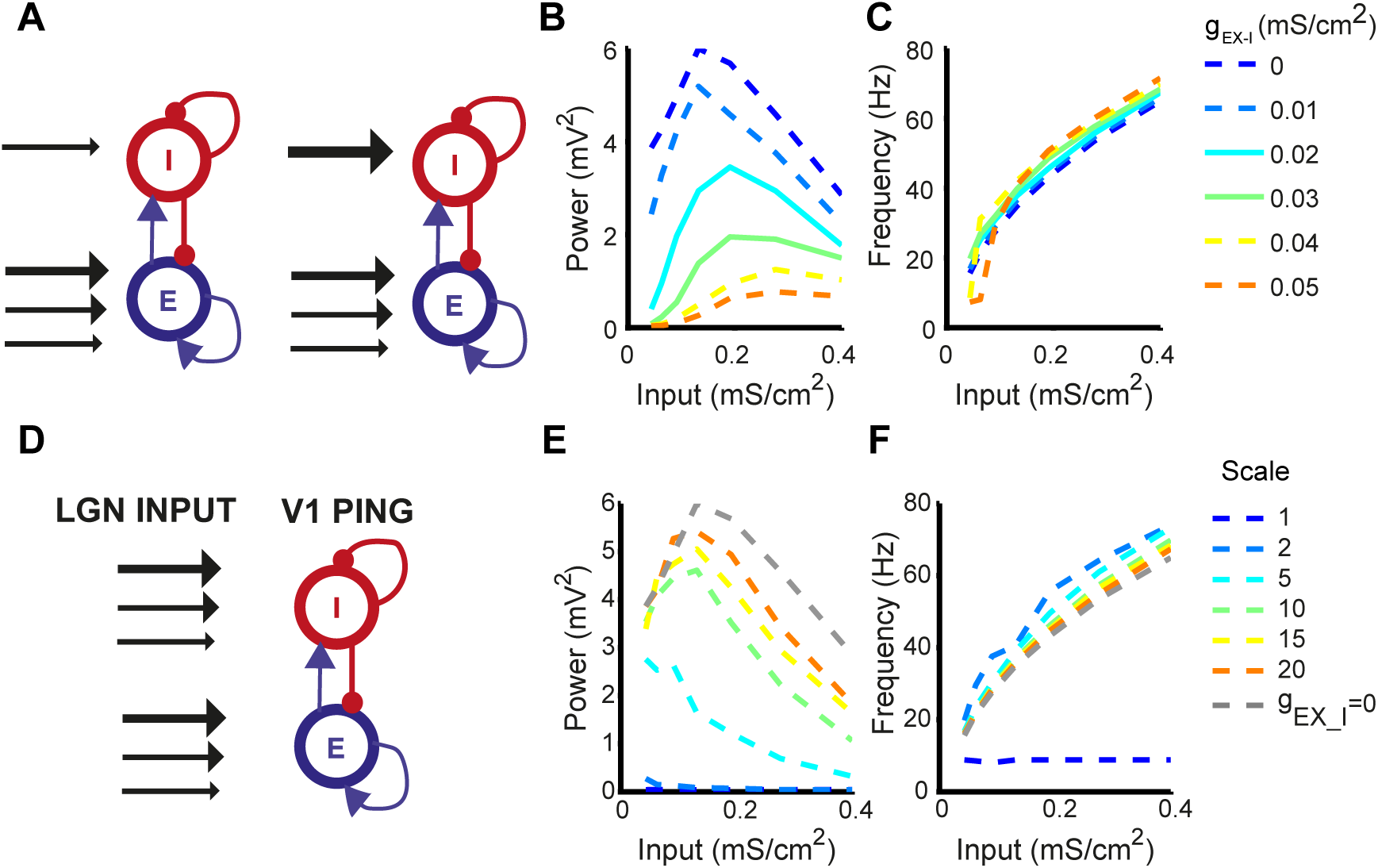
Investigation of effects of input to E and I populations: **A-C. Role of strength of fixed input to I cells and contrast modulated input to E cells. A**. Network diagram indicating manipulation of input to I cells, which was fixed across contrasts. Simulations were performed for different conductance magnitudes g_EX−I_, which influenced both the magnitude and heterogeneity of input to the I cells. Note that the conductance was fixed across all contrasts) as illustrated by the thickness of the arrow of input to the I cell population. As in previous simulations, the magnitude and heterogeneity of input to E cells varied with contrast (lines of different thickness). Results of the simulations in this scheme are shown in panels B and C. **B**. Peak power of the LFP as a function of contrast (input to the E cells). Different colours indicate the magnitude of the input to the I cells (parametrized by g_EX−I_). Continuous lines indicate valid and dashed lines invalid networks. Note that valid networks were only obtained at intermediate input magnitudes. **C**. Peak frequency of the LFP as a function of contrast (input to the E cells). Different colours indicate the magnitude of the input to the I cells (parametrized by g_E)−I_). **D-F. Effect of Contrast-modulated input on both E and I cells. D**. Network diagram indicating manipulation of input to I cells, which varied across contrasts as indicated by the changing thickness of the arrows. As in previous simulations, the magnitude and heterogeneity of input to E cells also varied with contrast (lines of different thickness). Colour-coded lines correspond to simulations for different scenarios each corresponding to a different scaling of the contrast-modulated input to the I cells relative to the input to the E cells, which was parameterized by Scale, so that 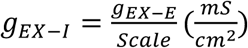. Results of the simulations in this scheme are shown in panels D and E. **E**. Peak power of the LFP as a function of contrast (input to the E cells)-same conventions as in B. **F**. Peak frequency of the LFP as a function of contrast (input to the E cells)-same conventions as in C. Note that when contrast-modulated input was varied simultaneously in both populations, a valid network was not be obtained even when the input to the I cells was up-scaled by a factor of 20 versus the input to the E cells (Scale = 20).

#### Contrast modulated input on both populations

We further investigated whether varying the input to *both* E and I cells across contrasts would result in valid network behaviour (see network diagram in **Figure 5D**). An important parameter here is the relative scaling of the input levels that E and I cells receive. We therefore performed simulations for different scenarios each corresponding to a different scaling of the contrast-modulated input to the I cells relative to the input to the E cells, which was parameterized as: 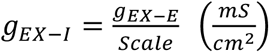. **Figure 5E** shows peak power and **Figure 5F** shows peak frequency as a function of the input strength to E cells with concomitant increases in input strength to I cells, for E-to-I ratio’s in input strength varying between 1 and 20. Throughout the tested conditions, we failed to obtain a valid network (i.e. at least one out of the five criteria was not fulfilled). These results show that the pace-making activity of I cells in weak PING is disrupted by a direct contrast-modulated input in a manner that leads to unrealistic model output. This confirms the crucial role of the precise levels of I cell desynchronization at high contrasts afforded by intermediate, fixed levels of input to the I cells – for generating power saturation or power reduction at high contrasts.

### 5. Oscillation destabilization at high contrasts is due to E cell mediated disruption of I cell pacemakers

Section 4 suggested that the state of de-synchronization in the I cell population controls the saturation or reduction of power at higher contrasts. In addition, Section 3 suggested that increased heterogeneity in activity of E cells as a function of input strength is required to produce power saturation or decay. Likely these two phenomena conspire to produce the power saturation or decay. It is possible that at high inputs, increased heterogeneity of E cells leads to a reduction of the I cell pace-making function and thus to network desynchronization. As a first approach to test this hypothesis, we selected the default valid, weak PING network and assessed its state of (de)synchronization as a function of input strength. The selected weak PING network exhibited fast power decay at high contrasts (**Figure 6A)** and generated realistic spiking outputs (**Figure 6B**). To assess the state of (de)synchronization within separate cell populations, we used three measures. First, we calculated the average maximum (across-lags) pairwise cross-correlation (MPC) between spike-trains of all model neurons. Second, we computed the phase locking value (PLV) between single unit spike trains and the LFP ((Berens, 2009; Lachaux, Rodriguez, Martinerie, & Varela, 1999), see Methods). Third, we calculated the coefficient of variation (CV2) value (Holt, Softky, Koch, & Douglas, 1996), which quantifies the variability of spike trains (see Methods).

**Figure 6:**
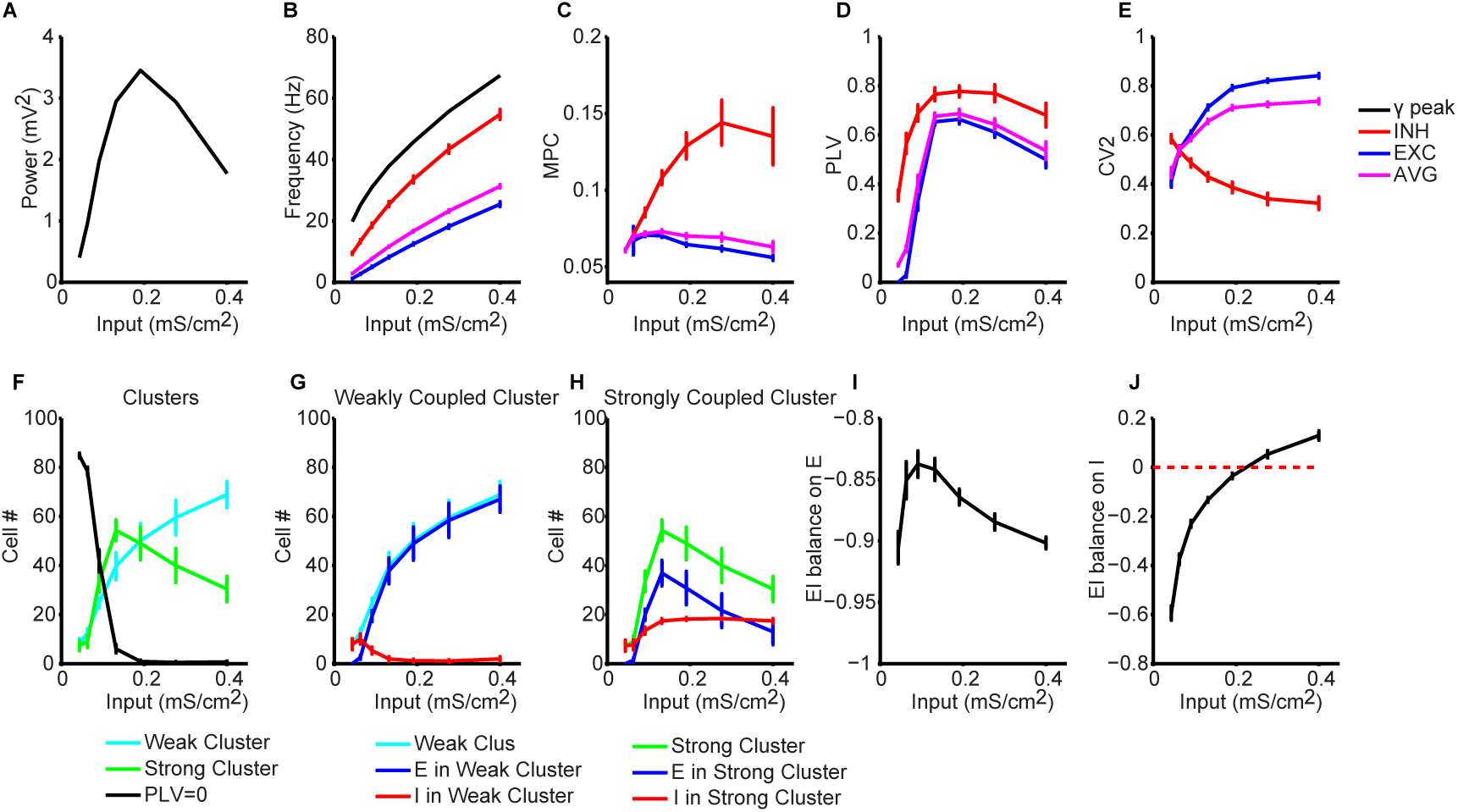
Investigation of Weak PING Power Decay. A-B Power and Frequency Analysis: The peak power **(A)** and the peak frequency **(B)** of the gamma oscillation as a function of contrast (input). In **(B)** superimposed on gamma peak frequency is the average firing rate of each cell population and of all the cells of the weak PING network (AVG). **C-E:** Spike train analysis. All measures are plotted as a function of contrast (input). The average maximum pairwise correlation (MPC) in panel **C**, phase locking value (PLV) in **D** and the coefficient of variation (CV2) in **E**, within each population and for all cells (AVG). **F-H:** Weak PING Clustering Analysis. On the left (**F**), the cells were classified separately for each contrast into two clusters based on their PLV value, indicated in cyan for the weakly coupled and in green for the strongly coupled clusters. The cells that were excluded from the clustering analysis due to low spike number are plotted in black. In **G** (weakly coupled cluster) and **H** (strongly coupled cluster), the participation of each cell type to the two clusters is illustrated with a blue line for E cells and a red line for I cells. **I-J:** E/I balance (EI_b_) as a function of input. EI_b_ is a proxy of the EI balance of the input that on average is received by each E cell **(I)** or I cell (**J**). It considers only the intra-network excitatory and inhibitory currents that each population receives (excluding the external stimulus-driven input).

The PLV (see **Figure 6D**) showed for E and I cells a similarly steep increase as a function of input up to intermediate inputs, which was then followed by saturation and decline. The observed change in PLV was qualitatively similar to the power variation as a function of input. Note that I cells were locked to the LFP more strongly than E cells. Further, we found that for I cells, MPC increased as a function of input strength before saturating at intermediate inputs (**Figure 6C**) while CV2 monotonically decreased (**Figure 6E**). Hence, I cell firing became more correlated and less variable as a function of input, whereby at high inputs a desynchronizing trend was observed (slight decline in MPC and decrease in PLV). By contrast, for E cells, the MPC, after reaching an initial low value at low inputs, tended to decline as a function of input strength (**Figure 6D**), while the CV2 was high and increased even more with input strength (**Figure 6E**). So, E cells were generally not well correlated in their firing and became more variable with increasing input levels.

The outcome of this analysis raises the question of how to reconcile the on one hand more regularly firing and better mutually correlated I cells at high input levels, with the decorrelation and increased variability among E cells on the other. To understand this, one should take into account that E cells in this setting experience two opposite forces. On the one hand, increased input to E cells will initially increase the drive to the I cells, which can then assert a pace-maker influence over the E cells. On the other hand, increased input also increases input heterogeneity, which acts against the pacemaker role of the I cells. Hence, the inherent increase of input heterogeneity with increased input levels may compromise the pacemaker function of the I cells. We surmised this is what may happen at high input levels, where the heterogeneity of the input to the E cells is highest and can no longer be mitigated by the pace-making (synchronizing) force of the I cells.

For this idea to be correct, it would have to be implemented in a subset of smaller clusters (or a single subcluster) of the network, in a manner that is obscured by the average behaviour of the network displayed in **Figure 6B**. It has indeed been reported before that in weak PING clustering, especially of E cells can occur (Kilpatrick & Ermentrout, 2011). It is therefore possible that increasing input heterogeneity may also split the population into clusters whose spikes are better coupled to the population LFP (strong cluster) than others (weak cluster). Thus, the behaviour of these clusters may yield more insight into the mechanism of power decay than the behaviour of I and E populations as a whole. A test of the clustering idea (**Figure 6F**) showed that beyond a minimum of input strength, a weak and a strong cluster could be distinguished, and that the membership of cells to the weak cluster increased with increasing input strength at the expense of membership to the strong cluster. This, in turn, could be related to the observed spectral power decline.

We then considered how the membership numbers of E and I cells to weak and strong clusters evolved as a function of input strength. A slight input increase from zero quickly moved all inhibitory cells from weak to strong clusters, in line with their pacemaker function in PING models (compare red lines in **Figure 6G and H**). The same slight initial input increase (until 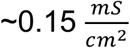) caused a similar increase of E cell membership to weak and strong clusters (similar initial increase of blue lines in **Figure 6G and H**). However, further increases in input (beyond 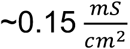) led to a fast increase of E cell membership to weak clusters relative to strong clusters. This confirms that I cells, despite their membership in the strong cluster (consistent with weak PING), at least partially lose their ability to set the pace of E cell spiking. This is in line with the slight decrease in MPC and PLV in I cells beyond input levels of 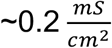 (**Figures 6Ba and 6Bb**).

To test further the mechanisms by which I cells would partially lose their pace making ability at higher input strengths, we looked at the balance of excitatory and inhibitory currents received by E and I cells (**Figure 6I-J**). The excitatory and inhibitory currents from all afferent synapses within the network were recorded from each cell and averaged across all cells, across time and trials yielding the average quantities *EI*_*b*_ for each cell population for the E and I cell group respectively (see Methods). E cells (**Figure 6I**) were dominated by inhibition throughout all input levels. Nevertheless, there was E a slight increase of E current with increasing inputs up to 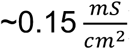), after which E currents declined again at higher inputs. In I cells (**Figure 6J**), the excitatory/inhibitory balance of inputs showed a monotonic increase as a function of input strength, during which a dominance of inhibition transitioned into a dominance of excitation at input levels of 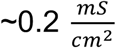 and beyond (black curve exceeding red dashed line). Hence, while self-inhibition dominated up to input levels of 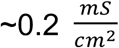 beyond this input level, noisy excitation dominated. That the I cells at high input levels became dominated by noisy currents of the E cells is remarkable, as it is the reverse of the standard situation in which I cells drive and pace the E cells. These observations are therefore relevant to understand oscillation destabilization at high input levels,

While the provided analyses do not pinpoint the exact mechanisms of power decay, they do constitute converging lines of evidence to support our hypothesis. In short, our analyses indicate that strongly and noisily driven E-cells partly compromise I cell pace-making function. This in turn results in power decay at high inputs. We should note that the reduction of I cell pace-making function was instigated by an increase in weakly coupled E cells (**Figure 6C**). Thus, in the validated model investigated here (**Figure 6A**), it is likely that E and I populations both contributed to oscillation destabilization.

### 6. GABA Differentially Modulates Gamma Power and Frequency in the weak PING model

Having a plausible generative model for gamma oscillations can be instrumental in addressing open questions that so far have been difficult to address in biological systems. It is well known that gamma oscillations and their spectral characteristics such as peak frequency and power depend on GABA-ergic inhibition (Brunel & Wang, 2003). A number of studies have reported that increased GABA conductance increases the power and decreases the frequency of spontaneous gamma oscillations (Faulkner, Traub, & Whittington, 1998; Lozano-Soldevilla, ter Huurne, Cools, & Jensen, 2014; Oke et al., 2010; Traub, Whittington, Colling, Buzsáki, & Jefferys, 1996; Whittington, Jefferys, & Traub, 1996; Whittington, Traub, & Jefferys, 1995). However, other studies have reported an increase in gamma peak frequency with increased GABA observables (Kujala et al., 2015; Muthukumaraswamy, Edden, Jones, Swettenham, & Singh, 2009). These discrepancies in the literature may be due to the discrepancies of modulating and quantifying GABA-ergic effects but may also reflect differential action of gamma generating mechanisms. In this section, we used the default validated weak PING model described in section 2 (Figure 2D), to investigate the role of GABA conductance. To that aim, we increased the conductance of GABAergic synapses in our model by 25, 50 and 100% (compared to the *default* weak PING, see Table 3 in Methods) for synapses on E cells alone, on I cells alone, and on both cell populations (**Figure 7**).

**Figure 7:**
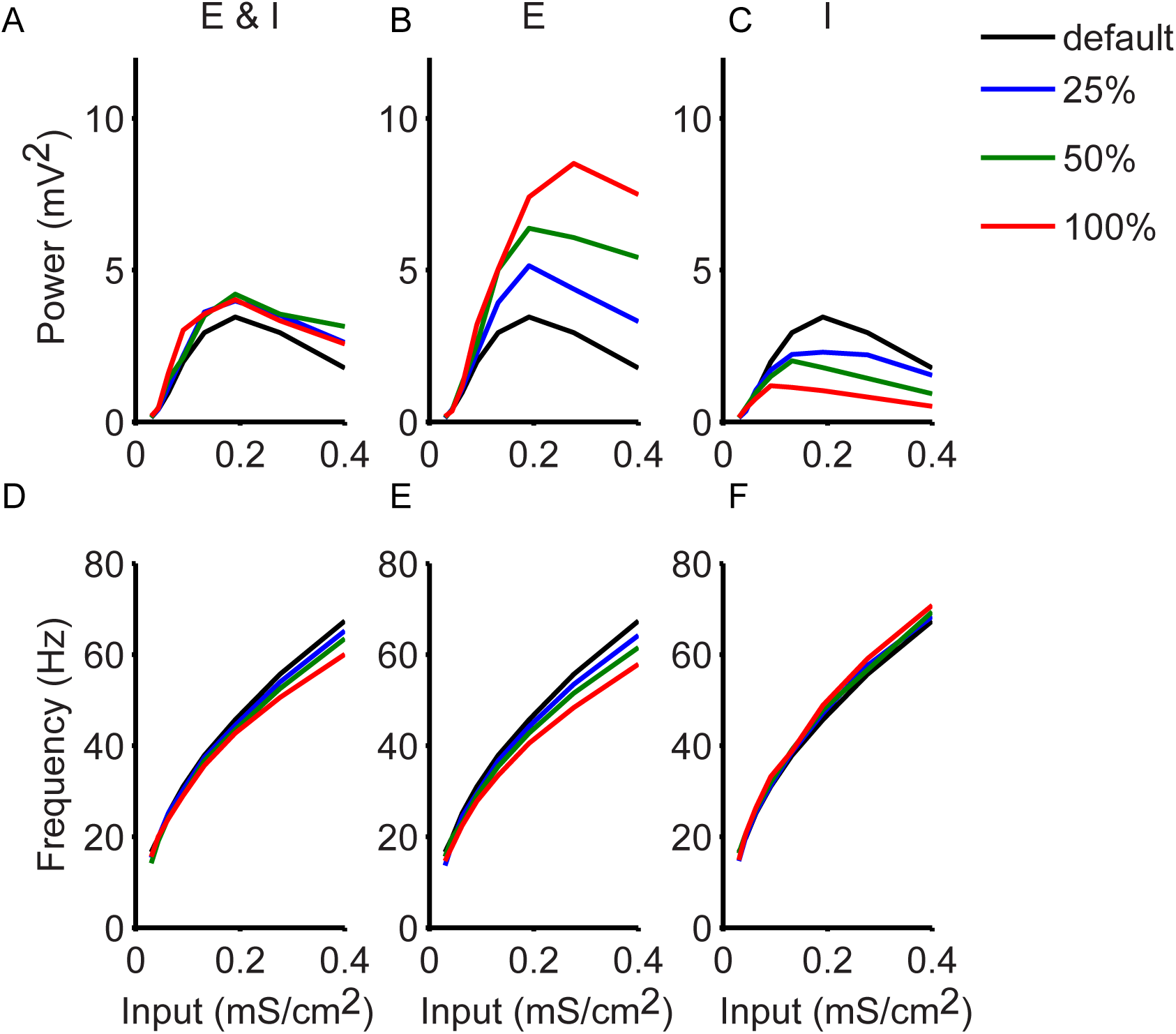
GABA Modulates Gamma Power and Frequency. Each column shows the results for increasing GABAergic conductance by 25, 50 and 100 % (colour-coded) on both E and I cells (**A and D**), only on E cells (**B and E**) and only on I cells (**C and F**). The results of our default weak PING are also plotted (black line). Top row shows the LFP power and bottom row the LFP frequency for the three cases. GABAergic enhancement only on E cells (**B and E**) resulted in the same but more pronounced effect as on both populations (**A and D**) whereas GABAergic enhancement only on I cells (**C and F**) reduced the power of oscillations and had a small increase effect on frequency.

When GABA conductance was increased above the default value in both populations, power increased (**Figure 7A**) and frequency decreased (**Figure 7D**), although this was not observed for low input strengths. GABAergic enhancement only to E cells resulted in an effect that was more pronounced but otherwise similar to the effect of GABAergic conductance enhancement to the E and I cells combined (**Figure 7B, E**). However, GABAergic enhancement only to I cells led to power reduction and a small frequency increase (**Figure 7C, F**). Overall, these results show that matching parametric empirical data against the pattern of model output in the different modelling scenarios can help in formulating informed hypotheses as to the biological mechanisms of empirically observed spectral effects of modulations in GABA conductance.

## DISCUSSION

We developed a set of five empirical criteria for assessing the validity of parameter settings in PING models. Empirical criteria were drawn from monkey V1 LFP and unit recordings obtained in a paradigm involving stimulus contrast variation (Roberts et al., 2013) supplemented by unpublished findings from our group, and experiments from (Contreras & Palmer, 2003). The empirical observables of recorded spikes and LFPs in response to gratings that varied in contrast were compared to model spikes and LFP proxy in response to input strengths at varying levels to E model units. This permitted an assessment of the match of the model’s behaviour with the empirical data, which would in turn indicate whether model parameter settings were valid or not. The match between model observables and empirical data was maximized by simultaneously setting a number of parameters so that all empirical criteria pertaining to both LFP and unit spiking responses were met simultaneously.

Our work led us to the following conclusions. First, in the paradigm we used, the gamma generating mechanism in primate (Macaque) V1 is more consistent with weak than with strong PING. Second, valid model behaviour for both the LFP and unit spiking requires specific interpopulation connection probabilities. Within weak PING, only certain subregions in EI/IE connection probability parameter space yielded valid models. Further, models showing power decay or power saturation at high input strengths were separable in this parameter space, thereby reproducing observed individual (Roberts et al., 2013) and interspecies variability (Hadjipapas et al., 2015; Self, Peters, Possel, & Reithler, 2016). Third, valid model behaviour poses highly specific requirements with respect to the input to E and I cells. Contrast modulation of input to the network needs to affect mainly the E cells, whereby its strength needs to scale with its heterogeneity. Input to the I cells should be moderate and should not vary with contrast. Fourth, using a validated weak PING model, we also discovered a possible route to partial oscillation destabilization, which we suggest underlies the LFP power saturation/decay at high contrasts. Specifically, an increase in input strength is accompanied by an increase in the heterogeneity of input to E cells, resulting in partial desynchronization of E cells. This is thought to disrupt the pace-making activity of the I–cells leading to partial oscillation destabilization. Finally, using the validated model, we found that, in order to replicate experimental results in terms of the frequency and power of gamma oscillations, GABA conductance input should be varied either for both E and I populations or only for the E population, but not the I population alone. Taken together, these findings facilitate the interpretation of existing and future experiments.

The findings in our work are relevant for understanding the mechanistic aspects of oscillatory neural responses and communication. Our approach of developing empirical constraints for gamma generation models is broadly applicable in existing datasets. For example, any dataset that records spikes and LFPs and includes a stimulus input manipulation (not limited to contrast) can be used to constrain gamma models using criteria similar to those used here. Such data are available in neurophysiological studies using rhesus monkeys (e.g., see (Berens, Keliris, Ecker, Logothetis, & Tolias, 2008; Frien, Eckhorn, Bauer, Woelbern, & Gabriel, 2000; Gieselmann & Thiele, 2008; Henrie & Shapley, 2005; Jia et al., 2013; Peter et al., 2019; Ray & Maunsell, 2010, 2011) or other species(Adesnik, 2018). It should be noted that, in datasets offering additional observables, the same approach to empirically constrain model parameters can be followed. These additional constraints will refine, confirm, or reject the parameter ranges we propose, all of which would be beneficial outcomes. The approach we suggest here is not meant to advocate the use of the specific constraints that we could derive from our data. Rather, our study is meant to advocate for the systematic use of empirical criteria to constrain parameter settings in computational models of gamma. One of the main aims in our study was to deliver a principled and general-purpose approach to bridge the empirical and computational literatures in the domain of gamma oscillations. Often, these literatures are somewhat disjointed, not least, because the large majority of computational studies of gamma and other oscillations set model parameters by convention. Here we improve upon this by restricting parameter setting to within empirically justifiable ranges.

In addition, our modelling approach can be extended to contribute to a better understanding of the origin of MEG and BOLD signals. Examples of MEG datasets that could be used for that purpose are from (Orekhova et al., 2020, 2018), who show gamma power saturation and decay as a function of stimulus velocity, and our own previous work. Better models of gamma may also help in better understanding the relationship between gamma, spiking and BOLD and can constitute the building blocks of models of the BOLD signal (Dahlqvist, Thomsen, Postnov, & Lauritzen, 2019; Takata et al., 2015).

Our work is relevant in the context of current literature that aims to establish whether gamma is generated by ING or PING models, and whether these models are ‘weak’ or ‘strong’. ING and PING models (both weak (Börgers et al., 2005; Börgers, Franzesi, LeBeau, Boyden, & Kopell, 2012; Lowet et al., 2015) and strong (Börgers & Kopell, 2005; Roberts et al., 2013) varieties) have been used to describe emerging oscillations throughout the brain (Buzsáki & Wang, 2012). Some optogenetic experiments showing that gamma could be induced more easily by stimulating I cells rather than E cells could be seen as favouring ING (Cardin et al., 2009). However, both macaque (Vinck et al., 2013) and cat (Moca et al., 2014) studies suggest PING rather than ING as the more likely mechanism for gamma rhythm generation in V1. This is supported by the observed phase-lead by a few milliseconds of excitatory cells to inhibitory cells which is indicative of PING-like mechanisms (Csicsvari, Jamieson, Wise, & Buzsáki, 2003; Hasenstaub et al., 2005; Tukker, Fuentealba, Hartwich, Somogyi, & Klausberger, 2007; Vinck et al., 2013). In the context of this research, the fact that observations from an expansive neurophysiological dataset in macaque V1 unambiguously point to weak PING is a relevant finding. In coming to that conclusion, the inclusion of LFP as well as unit spiking data was necessary. This confirms the necessity of considering single unit behaviour (micro-scale, spikes) as well as population behaviour (meso-scale, LFP) data and their inter-relationships in modelling. Doing so avoids special purpose models that capture some aspects of empirical data but remain unrealistic with respect to other aspects.

It has been proposed that weak PING is associated with attention and arousal, while strong PING is associated with active coding and formation of cell assemblies (see (Cannon et al., 2013) and references therein). In a study of (Börgers et al., 2005) that modelled the effects of attention in terms of PING circuitry, weak PING provided a background rhythm but stimulus representation occurred through (location) selective strong PING. We did not investigate attention here, and therefore cannot exclude that attention-induced stimulus representation occurs through strong PING. This in fact remains an interesting prediction that can be tested in a different empirical paradigm. Note that the neurophysiological dataset that we used to constrain our data was obtained by using responses to non-attended stimuli. This was true for the neurophysiological datasets we used from our own group (see Methods), as well for the study of (Contreras & Palmer, 2003). What can be therefore clearly inferred from our data is that stimulus processing of non-attended stimuli is associated with weak PING.

In our empirical data, some monkeys showed power decay and others power saturation at high contrasts. This individual variation occurred despite comparable experimental paradigms in the two monkeys. This suggests possible differences in the gamma generating networks among monkeys. In our modelling work, we found that only certain combinations of EI and IE connection probabilities yielded valid behaviour. Further, valid models exhibiting power saturation or power decay were separable in the parameter space spanned by these interpopulation connection probabilities. This supports the idea that some individual variability could be explained by differences in network connectivity. The inter-individual differences in network connectivity are likely strongly influenced by genetic factors (van Pelt, Boomsma, & Fries, 2012). In addition, although there is a remarkable evolutionary preservation of the different oscillatory brain rhythms (Buzsáki, Logothetis, & Singer, 2013), detailed aspects of gamma generating networks may differ between species. In contrast to LFP data in the awake monkey (Ray & Maunsell, 2010; Roberts et al., 2013)recent LFP gamma data recorded from awake human participants (Self et al., 2016) did not show power saturation with contrast. Thus, one could speculate that this could reflect a species difference in the specific network architecture generating gamma. Although significant differences exist in the methodology and scales of measurement, human MEG studies (Hadjipapas et al., 2015; Hall et al., 2005; Perry, Brindley, Muthukumaraswamy, Singh, & Hamandi, 2014) have also reported a lack of saturation at high contrasts, consistent with the human LFP data and therefore in contrast to the monkey LFP data. At this point, we must note, that the fact that gamma power saturation is not a general phenomenon observed in all experimental contexts should not be taken to mean that our modelling approach is also not generally applicable. Indeed, we show clearly that various degrees of power decline, saturation, or the absence thereof can all be understood within the context of our model. That is, a model that is ‘invalid’ for our own empirical data, may be ‘valid’ for other data. The underlying difference in parameter settings that are valid in different settings will be informative as to how gamma mechanisms differed among those studies.

In our data, gamma power showed a non-monotonic behaviour with contrast. At low contrasts, power increased and was maximal at middle contrasts. At high contrasts, power decayed or saturated. This suggests variation of synchronization with contrast and some degree of desynchronization (instability of the oscillation) at high contrasts. We investigated this idea in a single instantiation of a validated model. The hypothesized changes in synchronization were indeed visible in the model. In this model, the route of oscillation destabilization involved desynchronization of initially the E cells due to heterogeneous direct input. In turn this led to less synchronous recruitment of I cells partially breaking down the rhythmic synchronization. This mechanism of oscillation destabilization at strong input levels is different from other mechanisms described in a previous study (Börgers & Kopell, 2005). In that study, mechanisms by which PING rhythms were lost due to modulation of external input were investigated in a strong PING framework. Gamma rhythm became unstable for too strong input to I-cells (or too weak input to E cells) due to either “phase walkthrough” of I cells or “suppression of E cells”. In “phase walkthrough”, I cells synchronize but get ahead of the E cells (i.e., I cells fire without being prompted by E cells). In “suppression of E cells”, I cells fail to synchronise and fire asynchronously whereby their activity ultimately diminishes or ceases the E cells spiking activity (Börgers & Kopell, 2005). In our weak PING model, the second mechanism (“suppression of E cells”) was clearly not the case, as the activity of E cells was not suppressed when power decayed for higher inputs (see E cell average firing rate in **Figure 3Ba**). Hence, the mechanism which drove the synchronous rhythm by stimulating I cells was not abolished. The “phase walkthrough” entails I cells firing synchronously, but getting ahead of the E cells, i.e. spiking without being prompted by the E cells. This was also not the case in our weak PING model, as the I cells did not over-respond but rather failed to generate a coherent population response. Börgers and colleagues recently examined the effects of heterogeneity in synaptic strengths and in external drives to E and I cells in PING networks (Börgers et al., 2012). They found that, while the effects of heterogeneous input to I cells could be easily mitigated with strengthening of excitatory synapses, the effects of heterogeneous input to E cells remained pronounced even when inhibitory synaptic input to E cell was very strong. This is consistent with what was observed in our validated model, where synchronization broke down when heterogeneity in external input to E cells was too high. Börgers et al (Börgers et al., 2012) also predicted that breakdown of synchrony in a heterogeneous network can be predicted by the latency by which synchrony sets in, in a homogeneous network. A further prediction by Börgers et al. was that in a heterogeneous network a PING rhythm is established rapidly or not at all. In our setting it is difficult to test these predictions, as in weak PING synchronization is partial (noisy) both in terms of cell participation but also in time. Thus, synchronization varies over time, making an estimation of latency difficult. However, what is clear from our models is that weak PING can be obtained even in the setting of substantial heterogeneities in synaptic connection probabilities and direct inputs.

We have developed a model which is simplified enough to allow controlled manipulation of key parameters, such as the inhibitory and excitatory activity within the network, yet, detailed enough to reproduce our experimental findings and generate informed predictions on the underlying functional network circuitry. Despite the simplifications in our models, we kept a degree of realism by using conductance based rather than integrate-and-fire model neurons. It has been argued that biophysical detail in synaptic models is crucial for correctly simulating gamma dynamics (Cavallari, Panzeri, & Mazzoni, 2014), and using conductance-based models at least partially allays this concern. Nevertheless, there are limitations to our work. When modelling networks in visual cortex there are a number of important factors to consider such as using an appropriately large number of neurons (van Albada, Helias, & Diesmann, 2015), correctly modelling laminar structure and connectivity (Potjans & Diesmann, 2014) and taking into account different neuronal types and their morphology and thus appropriately constructing the LFP forward model (see (Einevoll, Kayser, Logothetis, & Panzeri, 2013) and references therein). Here we chose to validate a much-reduced point-neuron model in order to limit the number of parameters as much as possible to those that could be informed by the data. In addition, the model does not consider complex interactions with other brain processes, such as saccades (Martinez-Conde, Macknik, & Hubel, 2004; Martinez-Conde, Otero-Millan, & Macknik, 2013) or microsaccades (Lowet, Roberts, Bosman, Fries, & de Weerd, 2015) which influence gamma in-vivo. Moreover, our PING models were constrained by data collected across all cortical layers of V1. In the future, it will be interesting to subdivide the empirical data according to layers, to generate distinct PING networks for different layers. The work presented here shows how the model can be adjusted to match empirical data from two monkeys showing different power saturation characteristics. Likewise, different empirical constraints emerging from analysis of different layers should allow us to generate layer specific models.

The increased validation of local network models offered by our study can help advance the empirical validation of other models and can complement and inform hypothesis-driven experimental data analysis. Models like ours can be useful for deriving predictions for optogenetics experiments aiming to understand the network underpinnings of gamma signals, which is a fast growing new field (Cardin, 2016; Cardin et al., 2009; Hakim, Shamardani, & Adesnik, 2018; S. Sohal & Cardin, 2016; V. S. Sohal, Zhang, Yizhar, & Deisseroth, 2009; Veit, Hakim, Jadi, Sejnowski, & Adesnik, 2017). For example, the different factors that in our model are hypothesized to lead to power decay at high contrasts can be experimentally manipulated using optogenetics. In turn, optogenetics could identify separate roles of different classes of interneurons (see (Veit et al., 2017)), which could then be incorporated in the model. Moreover, modelling approaches as offered in the present study are relevant in the context of promising research suggesting that gamma entrainment by optogenetic tools and non-invasive brain stimulation can ameliorate deficits in for example Alzheimer’s disease (Adaikkan & Tsai, 2020; Iaccarino et al., 2016; Martorell et al., 2019) and schizophrenia (Cho et al., 2015; Cho & Sohal, n.d.; Gonzalez-Burgos, Cho, & Lewis, 2015). In particular, models as presented here may help to increase understanding of the mechanistic aspects of gamma generation that enable stable entrainment.

In conclusion, we suggest that insights from simplified models as used in the present study can form a basis for further constraints in more detailed and larger-scale biophysical models that consider neuronal morphology, laminar and horizontal structure and are set at an appropriate scale. In addition, the empirical validation approach presented here can be reiterated when upscaling such models. This is the subject of future work. Finally, we believe that the more general approach adopted here and elsewhere (see (Sherman et al., 2016) for tightly empirically-validated models of somatosensory beta rhythm), where experiment and theory go hand in hand and mutually reinforce each other is one that is much needed in systems neuroscience.

## Acknowledgments

Work was supported by European Union Seventh Framework Programme (FP7/2007-2013) under grant agreement no. 604102 (Human Brain Project) to AH, PDW and MR and NWO grants 016.105.203 and 453-04-002 to MR and PDW respectively.

## Author Contributions

MZ, AH conducted modelling and analysis, MR, PDW and EL collected and analysed empirical data. All authors contributed to conceptualising and writing the paper.

## METHODS

### Experimental Data Acquisition and Analysis

#### Acquisition and Analysis

We used existing datasets from two monkeys, the first (monkey S) being a dataset previously presented in Roberts et al., 2013 and the second (monkey O) being a similar dataset that has not previously presented. We choose to use this second dataset since, in the second monkey of (Roberts et al., 2013), peak gamma frequency was hard to estimate for low contrasts, where gamma overlapped with a pronounced beta band. However, note that our findings reported here were fully compatible with that data, and with additional data from monkey S recorded on the opposite hemisphere.

The monkeys were head-fixed and placed in a Faraday-isolated darkened box at a distance of 57cm from a computer screen. Stimuli were presented on a Samsung TFT screen (SyncMaster 940bf, 38°x30° 60Hz). The screen was calibrated to linearize luminance as function of RGB values. During stimulation and pre-stimulus time the monkey maintained eye position (measured by infra-red camera, Arrington 60Hz sampling rate) within a square window of 2×2°. This window was relatively large to allow for noise associated with the camera. In Monkey S the task was simply to maintain eye position within the window, in Monkey O the task was to maintain fixation until a colour change occurred at the fixation spot and reward was given for reporting the colour change by making an eye movement towards a target presented at the top of the screen. Stimuli were circular patches of square-wave grating with a spatial frequency of 2 cycles/degree. Contrast values in Monkey S were: 2.5 3.7 6.1 9.7 16.3 35.9 50.3 72% and 3 5 8 14 24 40 67 100% in Monkey O. Contrast values were chosen to keep the gratings, as much as possible, isoluminant with the background for all contrasts, with equal contrast between the background and both the black and white stripes. In Monkey S stimulus diameter was 5°, but varied in some sessions between 1° and 9°. In Monkey O stimulus diameter was 5° in all sessions. Eccentricity was between 5° and 6° in both monkeys.

Monkeys were implanted with recording chambers above early visual cortex, one positioned over V1/V2 and a second over V4. For the experiment reported here we used data from the V1/V2 chamber only. A head post was implanted to head-fix the monkey during the experiment. All the procedures were in accordance with the European council directive 2010/63/EU, the Dutch ‘experiments on animal acts’ (1997) and approved by the Radboud University ethical committee on experiments with animals (Dier Experimenten Commissie, DEC). Recordings were made with Plexon U-probes (Plexon Inc.) consisting of 16 contacts (10µm diameter, 0.5-1mW impedance, and 150µm inter-contact spacing). The probes were then advanced by a microdrive (Nan Instruments LTD.). The probes were connected to headstages of high input impedance, and data were acquired via the Plexon ‘Multichannel Acquisition system’ (MAP, Plexon Inc.). The measured extracellular signal was filtered online between 150Hz and 8kHz to extract spiking activity and filtered between 0.7Hz and 300Hz to obtain the LFP. The signal was amplified and digitized with 1kHz for the LFP and 40kHz for the spike signal. The data was converted from Plexon to Matlab file format and cut into trials from fixation onset to stimulus offset using the fieldtrip toolbox (Oostenveld, Fries, Maris, & Schoffelen, 2011). Line noise was removed from LFP using the fieldtrip dft filter, which fits a sine and a cosine at 50, 100 and 150Hz and subtracts these components from the data. Data was then further cut into non-overlapping 500 ms snippets excluding the first 500ms after fixation onset and the first 250ms after stimulus onset. LFP spectra were computed using a multi-taper method with discrete prolate spheroid sequences (DPSS) for frequencies 6 to 80Hz (smoothing ± 3Hz). Power spectra recorded in the stimulation period (S) were Z sored with respect to power spectra recording during the pre-stimulus baseline (B) computed over the 500ms prior to stimulus onset [Psi=(S-B)/standard deviation(B)]. The 1/frequency drop-off of power spectra was thus removed from the data.

#### Single unit spike sorting

Spikes were detected online by threshold crossings, where the threshold was set automatically and adjusted manually. Spike timing and shapes around the threshold crossing were recorded at 40Khz. To isolate single units, spike amplitudes were first normalized by their standard deviation and the shape was then decomposed into Haar wavelet features. The distribution of wavelet features was tested for normality and spike sorting was based on the non-normalised spike amplitude and the 10 wavelet features with the highest deviation from a normal distribution. We used Gaussian mixture models to perform final clustering. Starting with four initial clusters we reduced the starting value until the clustering converged. Following clustering we examined the spiking behaviour of each cluster. Clusters representing single units were identified as having a low probability (<5%) of inter-spike intervals less than 2ms paired with an average firing rate (over the course of the entire recording) of more than 5Hz. Additionally, we excluded units which were not responsive to the visual stimulus, which we identified as those which did not increase their response with increased contrast.

From the distribution of single unit mean spike waveforms we attempted to classify units as excitatory or inhibitory. Previous studies have shown a biphasic distribution of spike width where approximately 20% of units had narrow spikes, and were classified as presumed inhibitory, while the remaining units had broad spikes and were classified as presumed excitatory. We found no such distribution and therefore did not classify our neurons. We speculate that this failure may be due to the more diverse population of cell types in area V1, which includes for example spiny stellate cells – small excitatory cells which likely have intermediate width spike waveforms-. We note that there are also no other studies to (our knowledge) in macaque area V1 which have shown this distribution.

#### Feature Selection

We extracted features from the single unit and LFP activity in order to validate our PING model. Results from single unit and the LFP activity were combined across all depths and we focused our analysis on the sustained period (300 ms post-stimulus onset). We also excluded from the analysis contrast values for which the highest frequency oscillations observed were less than 15 Hz (outside the gamma range of interest and overlapping with the beta band which was present for all contrast conditions). We took the same approach with the model for which we also ignore non-gamma frequencies. We observed that the model also starts from beta-like oscillations for low input values. Thus, the model and data suggest that gamma band activity may operate over a wide continuum of frequencies bands as a result of contrast modulation. Activity in vivo within low frequency ranges during low contrast stimulation reflects some combination of gamma band generators and beta band generators.

### Network Model

We modelled the response of V1 LFP and single unit activity during contrast modulation (during passive viewing of stimulus) 300 ms post-stimulus presentation. We built a spiking network model in order to validate our model’s different modalities (LFP, spiking activity) and used the validated model to gain to insight to the underlying mechanism of the gamma oscillation in V1 as modulated by contrast.

A summary of the model description according to the scheme proposed by (Nordlie, Gewaltig, & Plesser, 2009) is given in **Table 1**. The spatially unstructured model was comprised of two kinds of Hodgkin-Huxley type single compartment neuron models for the excitatory and inhibitory populations as in (Buia & Tiesinga, 2006). The I cells were modelled with the Wang-Buzsaki model (Wang & Buzsáki, 1996) developed for fast-spiking hippocampal basket cells. The E cells were modelled with the Golomb-Amitai model (Golomb & Amitai, 1997) model for regular spiking E cells. Note that the E cell model had a lower intrinsic frequency of the response to input (f-I curve) than that of the I cell model. The cells from the two classes were connected randomly (no spatial structure) in a sparse way with excitatory (AMPA) and inhibitory (GABA) voltage dependent synapses and 1ms delay (for detailed description see **Table 1** and **Table 2** and for illustration the network diagram in **Figure 2A**).

The contrast-modulated input from the LGN was described with a phenomenological model. Both classes received input in the form of Poisson spike train with fixed rate (yet different for each cell). When contrast was varied the input to the E cells was modulated (as per conductance magnitude – *g*_*EX*−*E*_) representing the LGN contrast-dependent input. Thus, we considered the frequency of the input as fixed and increased the conductance as the sum of many simultaneous spikes that make postsynaptic firing more likely due to the larger amplitude post-synaptic potential. A degree of heterogeneity was introduced to the network by means of its input frequency, which was drawn from a normal distribution (200 ± 25 Hz). Also, the cells were given an initial membrane potential drawn from a uniform distribution between [−90,50] mV for E cells and [−85, −45] mV for I cells (i.e. reversal potential for leak current ±20 mV).

To mimic the experimental procedure of averaging across trials, we initialized the network with a new random seed (which determined the within network connectivity and initialisation and the input variability) and ran it for all contrasts. We repeated this process for 10 trials and averaged across trials both for LFP and spike measures. In all Figures we report mean and 1 SEM values.

In order to extract the LFP signal we regarded the average of the inverse somatic membrane potential of the E cells as a mean field signal (see **Table 1F**). An alternative was to use the sum of absolute values of AMPA and GABA currents, as in (Mazzoni et al., 2008). We implemented both measures and generated similar results for LFP power and frequency (not shown here). A recent study (Mazzoni et al., 2015) showed that in integrate and fire network models, the membrane potential LFP was not the optimal measure therefore other measures of weighted combinations of synaptic currents were proposed. However, the spectral peak and frequency characteristics were retained across measures (see figure 5 in (Mazzoni et al., 2015)). In our conductance-based network we opted to use the membrane potential LFP proxy rather than the synaptic current proxy in order to compare directly with other models which used a current–based rather than a synaptic-based contrast input (e.g. the strong PING from (Roberts et al., 2013)).

### LFP model

The simulated LFP of the spiking network model was calculated as the average of the additive inverse somatic membrane potential of the pyramidal cells reflecting a mean field signal approximation of the activity of the population of neurons, as in previous modelling studies (Bazhenov et al., 2001; Bush & Sejnowski, 1996).

### Simulation Protocols

***Section 2:*** For **Figure 2** the weak PING model parameters are listed in **Table 1** and **Table 3**. Connection probabilities for EI and IE were varied in the range of [0.1,0.9]. The excitatory input to the network (i.e. contrast) was encoded through the variation of the synaptic conductance in the range 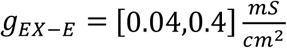. For **Figure 2C-E** the EI and IE connection probabilities were 2**C**: (0.1, 01), **2D**: (0.6, 0.7), **2E**: (0.7, 0.3). The weak PING parameters with the EI and IE connection probabilities fixed to 0.6 and 0.7 respectively is defined as the *default* weak PING which is used throughout the rest of the sections.

***Section 4:*** For **Figure 3A** the strong PING network model parameters and current-based input are described in **Table 3** and **Table 4** respectively. The *default* weak PING in terms of connectivity as defined in section 2 is used in **Figure 3B-D**. The conductance-based input for **Figure 3B** was the same as in **Section 2** (see **Table 4** - second column). The current-based input to weak PING in **Figure 3C** and **3D** are also described in **Table 4** – third and fourth column respectively. In **Figure 4A** different experimental scenarios are ran for input standard deviation 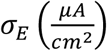 in the 0.01-3 range. Here the contrast is encoded as a function of input strength 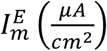 in the [0.43,8.0] range. In **Figure 4B** different experimental scenarios were ran for input with frequency variability of the poissonian train *F*_*S*_ in the range [0,250] Hz. Here the contrast was encoded by the increase the synaptic conductance in the range 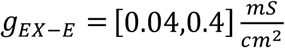.

***Section 5:*** In **Figure 5** the network is the default weak PING model with conductance-based input as in section 2. In **Figure 5A-C** the contrast-modulated input to I cells is fixed yet has a different magnitude per simulation scenario. This is parameterized by the synaptic conductance of the input to I cells *g*_*EX*−*I*_ which takes values in the range 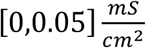. In **Figure 5D-F** the contrast-modulated input to I cells is the same as for E cells with a scaling applied in each simulation scenario parameterized as: 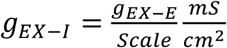 with *Scale* taking values in the range [*b*,20].

***Section 6*:** For **Figure 6 and S3** the network is the default weak PING model with conductance-based input as in Section 2.

***Section 7:*** For **Figure 7** the network is the default weak PING model as in Section 2. The GABA conductance in modified for each simulation scenario on either E cells 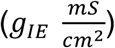 or I cells 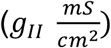 or both populations. The conductance in each case is increased by 25%, 50% and 100%.

## Supplementary

For **Figure S1** and **S3** the default weak PING model was used. For **Figure S2** the network was similar to the default weak PING model with conductance-based input, as in Section 2, with the network was upscaled by a factor of 10 from N=100 to N=1000 and the probabilities of connection divided by the same factor.

### Network Model Validation

From our experimental data set based on average responses (across 45 sessions in Monkey S and 53 sessions in monkey O) in two monkeys. From the literature we established empirical criteria robustly describing the gamma response over a range of contrasts. For a network to be considered valid it had to fulfil all these criteria simultaneously. The criteria are listed in **Table 5** and illustrated in **Figure S1**. We varied crucial parameters, i.e. the probabilities of connection between E and I populations systematically to produce model outputs (model LFP and spikes) for each parameter setting. The model outputs then needed to satisfy all the empirical criteria derived by the experimental LFP and spike data. Following the validation process, valid networks were labelled as “decay” or “saturation” networks by testing whether the contrast-modulated peak power decayed rapidly (decay) or slowly (saturation) using as a threshold for the decay slope the value 0.03 (unitless).

### Model Data Analysis

For all measures described in this section, the mean and SEM per population (E and I cells) was evaluated across the N=10 trials.

#### Variability

As a measure of spike-train variability we use the coefficient of variation (CV2) as proposed by (Holt et al., 1996). To obtain CV2 for a spike from a spiketrain with spike times *t*_*i*_(0 ≤ *i* ≤ *N*) we calculate the standard deviation of two adjacent interspike intervals (where *Δt*_*i*_ = *t*_*i*_ − *t*_*i*−1_), divide by their mean and multiply by √2, so that CV2 will have a mean of 1 for a Poisson process,

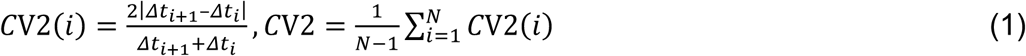

The average of CV2 over *i* gives the measure of variability of a spike train. Thus, compared to standard coefficient of variation (CV), CV2 provides a more reliable measure of intrinsic variability of spiking processes independent of slow gradual changes in firing rate. A Poisson process has a CV2 value of 1.

#### Correlation

We estimate the overall correlation of the network by considering the maximum pairwise correlation coefficient (MPC). That is the maximum absolute cross correlation across lags which was computed for each pair and then averaged across all pairs for each population. The MPC was computed for spike trains after subtracting the mean firing rate in MATLAB with xcorr function (maxlag=60 msec, option ‘coeff’). Note that we only considered cells that spiked more than 20 spikes during a trial.

#### Spectral Analysis

We estimated the Power Spectral Density (PSD) via the Thomson multitaper method in MATLAB (pmtm, with default settings) of the LFP (with the mean subtracted) per trial and then average across trials. The maximum peak power and respective peak frequency (frequency of peak power) per contrast were then extracted from the average spectral response.

#### Phase Locking

We computed the PLV (Lachaux et al., 1999) per each neuron using the CircStat MATLAB Toolbox described in (Berens, 2009). We collected the phases at which a neuron fired and from that phase distribution, we estimated the PLV. As in the MPC calculation, we only considered cells that spiked more than 20 spikes within a trial. To compute the PLV we filtered the LFP by the peak frequency per contrast ±8Hz after subtracting the mean. The PLV was computed from the whole trial for each cell and then averaged across all (1) I cells, (2) E cells and (3) across the whole population. Note that in averaging we included the cells for which a 0 value was assigned.

#### E/I Balance

The excitatory and inhibitory currents from all afferent synapses within the network were recorded from each cell and averaged across all cells. We denote as IIE(t) and IEE(t) as the total inhibitory and excitatory currents respectively for the E cell population and as Iii(t) and IEi(t) the total inhibitory and excitatory currents respectively for the I cell population. For a given cell (E and I) the E/I balance *B*_*E*_(*t*) and *B*_*I*_(*t*) was estimated by calculating

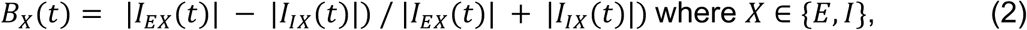

and then this was averaged across all E and I cells respectively and across time and trials yielding the average quantities *EI*_*b*_ for each cell population.

#### Clustering

The clustering was performed using custom Matlab, K-means algorithm with a pre-set number of clusters equal to 2, defining the group with the highest PLV values as the locked cluster and the group with the lowest PLV values as the unlocked cluster. K-means was selected to cluster the groups of cells based on their PLV response into two groups for each contrast in an unbiased way (i.e. without determining an a-priori cut off for each contrast point). Note that a value of PLV=0 was assigned to cells below the 20 spikes threshold to facilitate the clustering analysis detailed below yet their PLV values were not taken into account for the population PLV average. As a clustering feature we used the PLV value per cell and per trial (estimated for the trial duration) and the clustering algorithm was ran for each trial separately. The mean and SEM value of E and I cells per of their PLV values per cluster was evaluated across trials. To evaluate the validity of the clusters we calculate the mean and SEM silhouette value plotted in **Figure S3** for the points of each cluster across trials. The silhouette value is a measure of similarity for each point to points of its own cluster when compared to points in other clusters. It ranges from −1 to +1 and a high silhouette value indicates that a point is well-matched to its own cluster, and poorly matched to neighbouring clusters.

### Network Scaling

The network was upscaled by a factor of 10 from N=100 to N=1000 (**Figure S2**) by dividing the probabilities of connection by the same factor. Note that the conductances were normalised in all cases by the number of afferent connections, *c*_*XY*_ * *N*_*X*_, i.e. the product of probability of connection from X to Y population and the number of cells of the afferent population X.

### Simulation and Analysis

The simulation of the network was done in BRIAN 1.4.1 (http://briansimulator.org/) (Goodman & Brette, 2008).The equations were solved with Runge-Kutta solver with simulation timestep *dt* = 0.05*ms*. The results were analysed in MATLAB (MATLAB_R2014a - http://www.mathworks.com/)). The code is available upon request and will also be made freely available through the Human Brain Project repository.

## SUPPLEMENTARY FIGURES

**Figure S1:**
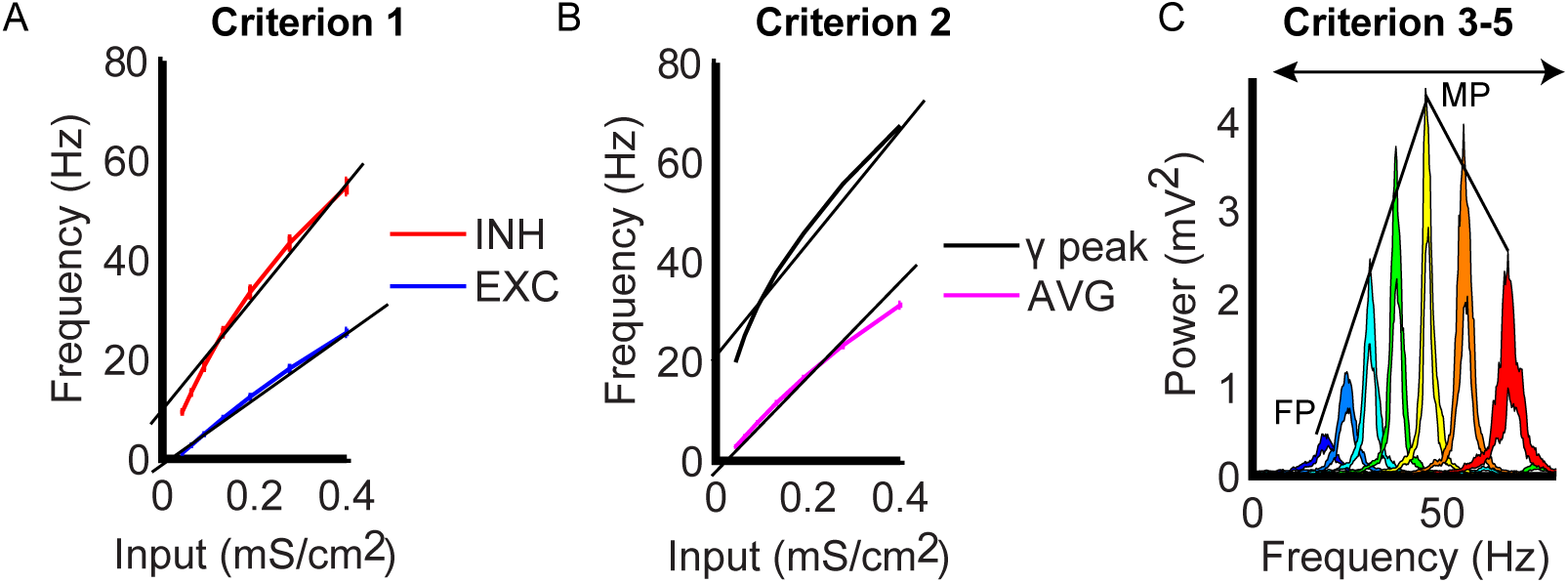
Network Feature Selection based on population firing rates and LFP spectra. **A:** The extraction of the slopes of the average firing rate of the I- and E-cell populations mean firing rate across all contrasts for each population. **B:** The extraction of the slopes of the average firing rate of the whole population and of the peak frequency of oscillations as well the overall average population firing rate and oscillation frequency. **C:** The extraction of features from the LFP spectra such as the first and the maximum power (FP and MP), the frequency range, the slope of the power rise and the slope of the power decay. For a detailed description see **Table 5**.

**Figure S2:**
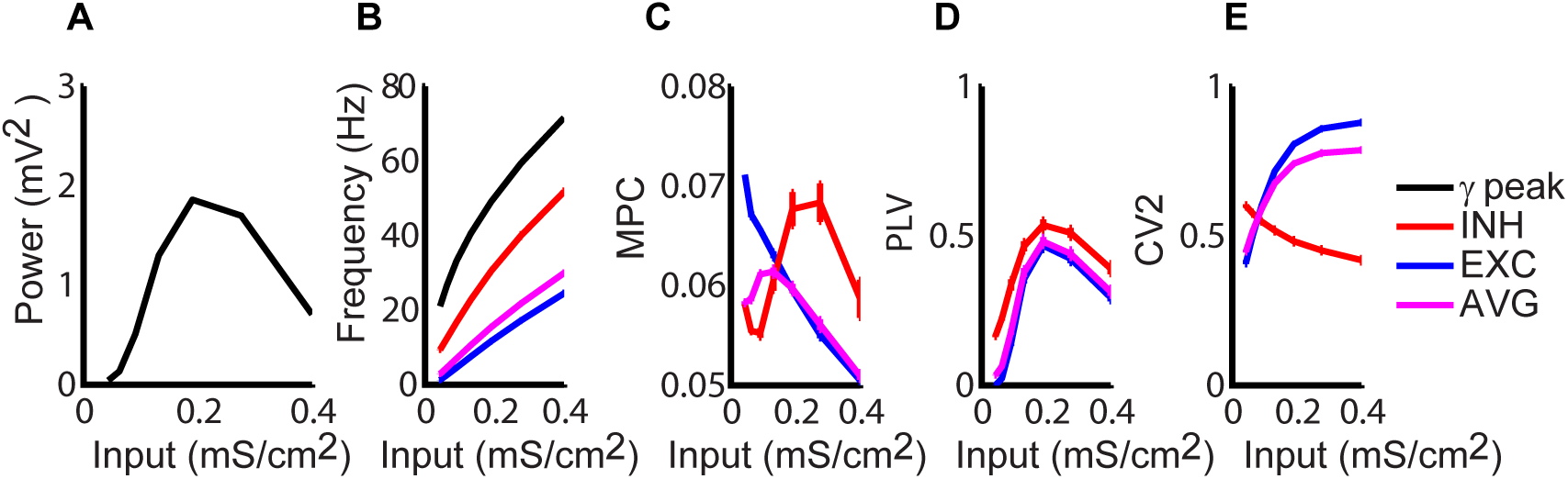
Weak PING with N=1000. An upscaled version of the default weak PING Network (see **Figure 6**) exhibiting similar results across both the spectral LFP and the single cell spiking behaviour. The conventions are the same as **Figure 6A-E**. The peak power per each contrast (first column) and the peak frequency of the gamma oscillation (second column) superimposed with the average firing rate of each cell population and of all the cells of the weak PING network. The average maximum pairwise correlation (MPC) on the third column, phase locking value (PLV) in the fourth column and the coefficient of variation (CV2) on the fifth column, within each population and for all cells.

**Figure S3:**
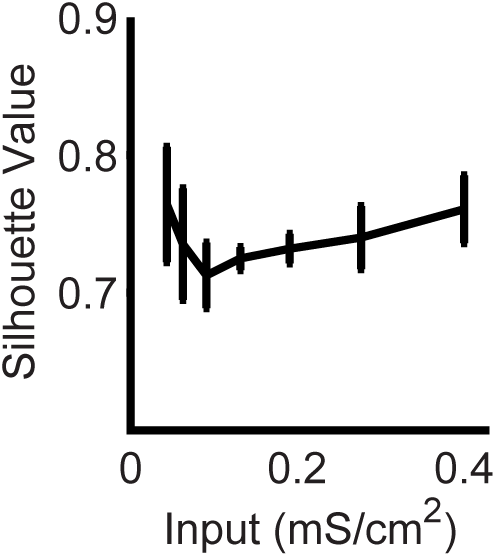
Cell Clustering based on PLV. Mean silhouette value of the clustering analysis based on PLV values. The silhouette value is a measure of how well clusters are separated. The measure ranges from −1 to 1, where 1 is maximal separation (for more information see Methods).

